# Mosquito cellular immunity at single-cell resolution

**DOI:** 10.1101/2020.04.08.032508

**Authors:** Gianmarco Raddi, Ana Beatriz F Barletta, Mirjana Efremova, Jose Luis Ramirez, Rafael Cantera, Sarah A. Teichmann, Carolina Barillas-Mury, Oliver Billker

**Affiliations:** Wellcome Sanger Institute, Hinxton, Cambridge CB10 2AZ, UK; Laboratory of Malaria and Vector Research, National Institute of Allergy and Infectious Diseases, National Institutes of Health, Rockville, MD 20852, USA; Crop Bioprotection Research Unit, United States Department of Agriculture, National Center for Agricultural Utilization Research, Agricultural Research Service, Peoria, IL, United States; Zoology Department, Stockholm University, Stockholm, S-10691, Sweden and Departamento de Biología del Neurodesarrollo, Instituto de Investigaciones Biológicas Clemente Estable, Montevideo 11600, Uruguay; Institute & Dept Physics, University of Cambridge, JJ Thomson Ave, Cambridge CB3 0HE, UK; Molecular Infection Medicine Sweden, Molecular Biology Department, Umeå University, Umeå S-90187, Sweden

## Abstract

Insect hemocytes are the functional equivalents of leukocytes and limit the capacity of mosquitoes to transmit human pathogens through phagocytosis, encapsulation, secretion of immune factors and immune priming (*1*, *2*). Here we profile the transcriptomes of 8506 hemocytes of *Anopheles gambiae* and *Aedes aegypti*, two important mosquito vectors. Blood feeding, infection with malaria parasites and other immune challenges reveal a previously unknown functional diversity of hemocytes, with different types of granulocytes expressing distinct and evolutionarily conserved subsets of effector genes. A new cell type, which we term megacyte, is defined in *Anopheles* by a unique transmembrane protein marker (TM7318) and high expression of LPS-Induced TNF-alpha transcription factor 3 (LL3). Knock-down experiments indicate that LL3 mediates hemocyte differentiation during immune priming. We identify two main hemocyte lineages and find evidence of proliferating granulocyte populations. We validate our analysis with RNA in-situ hybridization and highlight the mobilization of sessile hemocytes into circulation upon infection. Our data (https://hemocytes.cellgeni.sanger.ac.uk/) provide the first atlas of medically relevant invertebrate immune cells at single cell resolution. It provides an important resource for invertebrate immunology by identifying cellular events that underpin mosquito immunity to malaria infection.

## Main

*Anopheline* mosquitoes transmit *Plasmodium* parasites to humans, and are responsible for an estimated 219 million cases of malaria, leading to over 400,000 deaths annually(*3*). Parasites taken up by female mosquitoes from the blood of an infected human transform into motile ookinetes, which traverse the mosquito midgut and establish an infection. The mosquito’s immune system limits *Plasmodium* infection in several ways (*4*, *5*), and hemocytes, the insect white blood cells, are key players in these defense responses (*6*, *7*). Ookinete invasion triggers a strong nitration response in invaded midgut epithelial cells and their basal lamina (*8*, *9*). Hemocytes that come in contact with a nitrated midgut basal lamina release microvesicles into the epithelial basal labyrinth and promote local complement activation, inducing parasite lysis (*7*). An infection with *Plasmodium* primes mosquitoes to mount a stronger immune response to subsequent infections (*2*). Primed mosquitoes release hemocyte differentiation factor (HDF) into the hemolymph (*2*), consisting of a complex of lipoxin A4 bound to evokin, a lipocalin carrier(*10*). HDF increases the proportion of circulating hemocytes of the granulocyte type (Rodrigues et al., Sci 2010), promotes microvesicle release, and enhances complement-mediated parasite lysis (*7*). Enhanced immunity is lost if HDF synthesis is blocked(*10*).

Three hemocyte types have been described in *Anopheles gambiae* based on their morphology (*11*). Granulocytes are highly phagocytic cells of about 10-20 μm, while oenocytoids are relatively smaller (8-12 μm), round cells that produce melanin, an insoluble pigment involved in wound healing and pathogen containment by encapsulation. Finally, prohemocytes are small round cells (4-6 μm) with a high nuclear to cytoplasmic ratio, thought to be precursors of the other two cell types. Hemocytes can be circulating or sessile, and alternate between these two states (*12*, *13*). However, the full functional diversity of mosquito hemocytes and their developmental trajectories have not been established, and it is not clear to what extent morphologically similar hemocytes are functionally equivalent. Here, we use single cell RNA sequencing (scRNA-seq) to analyze the transcriptional profiles of individual mosquito hemocytes in response to blood feeding or infection with *Plasmodium*. We identify two basic lineages and differentiation pathways in prohemocytes and granulocytes, and we discover new hemocyte populations and markers of immune activation. We use *in situ* hybridization to integrate the transcriptional profiles with morphological analysis of circulating hemocytes. Newly identified markers reveal the dynamic behavior of sessile hemocytes in response to *Plasmodium* infection. A comparison of hemocyte types from *A. gambiae* and *A. aegypti* show that some of them are shared, while others appear to be unique to each mosquito species.

### scRNAseq reveals new granulocyte types and states

Circulating hemocytes were collected from adult *A. gambiae* M form (*A. coluzzii*) females that were either kept on a sugar meal or fed on a healthy or a *Plasmodium berghei*-infected mouse (Fig. 1a). Transcriptomes from 5,383 cells (collected 1, 3, and 7 days after feeding) revealed nine major cell clusters (Fig. 1b, Fig. S1). Two clusters originate from adipose tissue (fat body, FBC 1 and 2) and one from muscle tissue (MusC; Fig. 1b-c). FBC1 cells express several immune-modulatory genes such as CLIPs (CLIPA1, 7, 8, 9, 14), LRIMs (LRIM 1, 4A, 8A, 8B, 9, 17), lectins (CTL 4, MA2) and SRPN2 (Fig. 1b-c and Fig. S2, Table S1), while FBC2 cells express high levels of vitellogenin after blood feeding, a canonical marker of fat body(*14*). Based on their unique transcriptional profiles, we identified six hemocyte clusters (HC) (Fig. 1b, c). Hemocyte cluster 1 (HC1) has high mRNAs levels of prophenoloxidases, including PPO4 (Figs. 1b-c) and PPO9, characteristic of oenocytoids. This expression profile agrees with previously reported scRNA-seq data for a small number of hemocytes (25 cells (*15*)). To select markers for major hemocyte lineages, we used bulk RNAseq data from different tissues to guide us towards hemocyte specific genes (Fig. S3 and Table S2). HC1 contains low levels of leucine-repeat protein 8 (LRR8) mRNA, while HC2-4 have an inverse pattern (i.e. low or absent PPO4 and high LLR8 levels; Fig. 1b-c and 2a, Fig. S2).

**Fig. 1:**
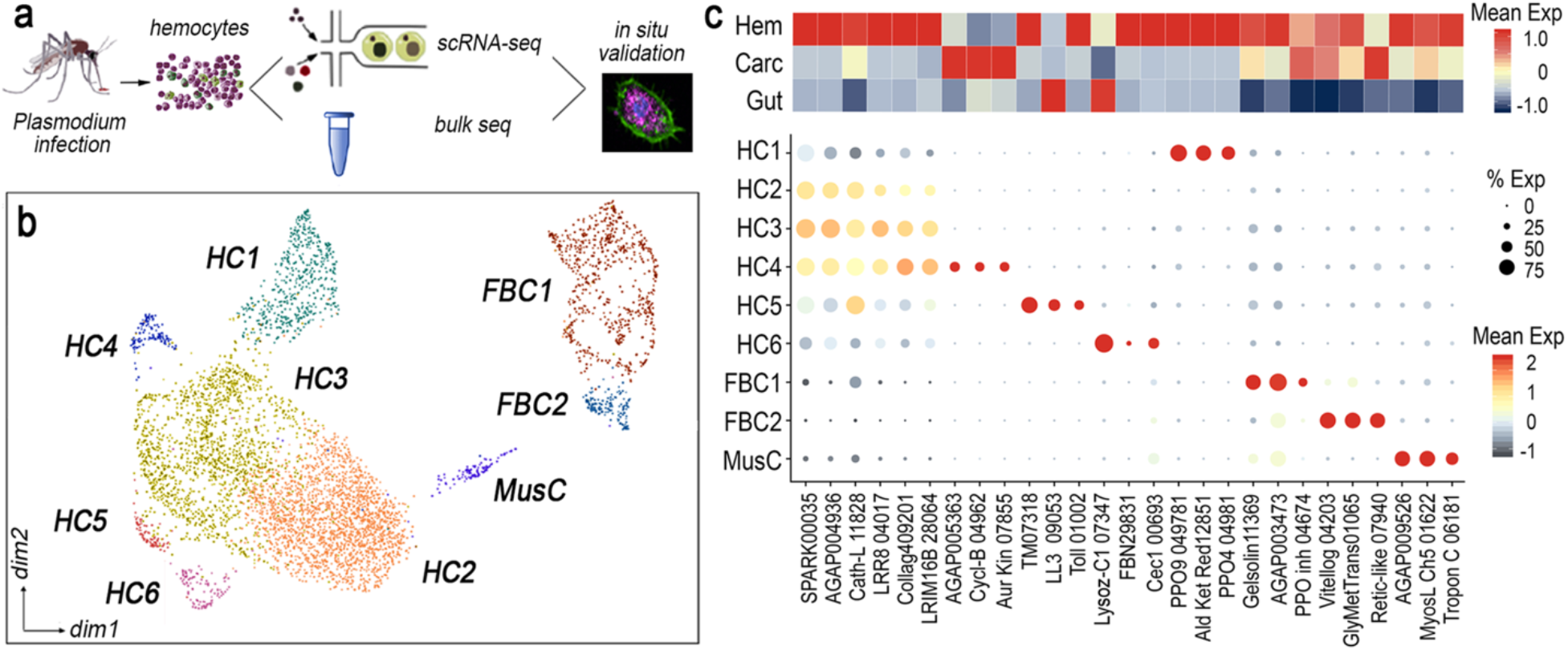
Molecular identification of hemocyte cell types of *Anopheles* mosquitoes. **a**, Workflow and experimental scheme. **b**, UMAP of 5383 hemolymph *A. gambiae* (M-form) cells colored by cluster (cell type) identity with Seurat clustering. HC, hemocyte cluster; Mus, muscle; FBC, fat body cluster. **c**, Top panel: heatmap showing mean expression of three gene markers per scRNA-seq cluster in bulk-RNAseq of mosquito tissues. Hem., hemocytes; Carc., carcasses; Gut, gut samples. Bottom panel: dot plot showing corresponding expression of the cluster markers genes, where color indicates mean expression and dot size encodes percentage of cells within the cluster expressing the marker. Last five digits of each marker gene are the Vectorbase transcript accession numbers (after AGAP0).

*In situ* hybridization using these markers (Fig. 1c and Table S3) revealed that the morphology of circulating HC1 (PPO4^high^/LLR8^low^) cells is typical of oenocytoids, with round cells that have few pseudopodia and granules (Fig. 2b); while the morphology of HC2-4 (PPO4^low^/LLR8^high^) cells is typical of prohemocytes and granulocytes (Fig. 2b). HC2 and HC3 shared many markers such as SPARC, cathepsin-L and LRR8 (Fig. 1c). However, HC2 had 73% fewer unique molecular identifiers (UMI) (mean UMI of 413) in comparison to HC3 (mean UMI of 1,516), suggesting that HC2 cells are less differentiated cells along this lineage, most probably prohemocytes and correspond to the observed small round cells with high nuclear to cytoplasm ratio (Fig. 2b), while HC3 cells have a typical granulocyte morphology, with prominent pseudopodia and abundant granules (Fig. 2b).

**Fig. 2:**
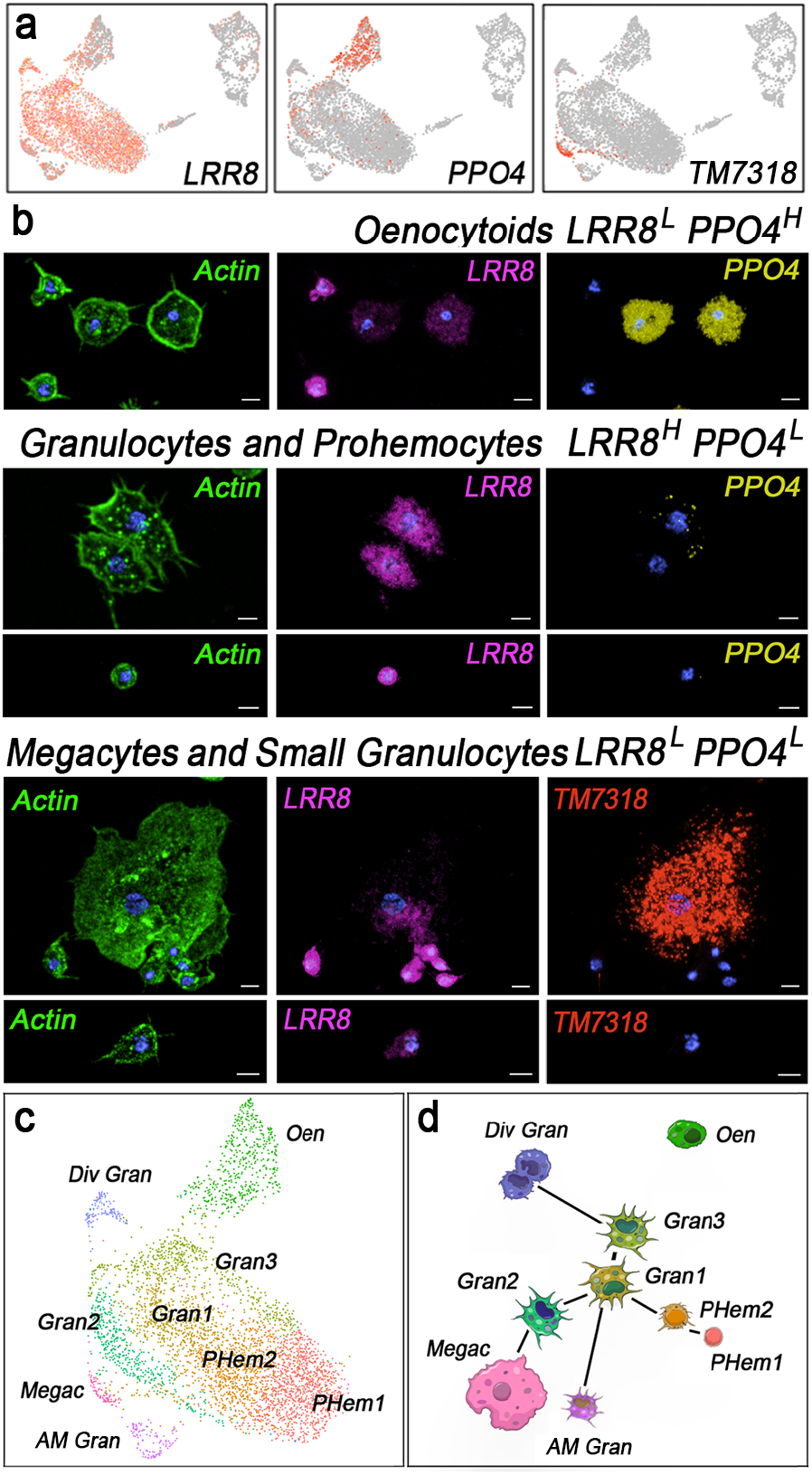
Cell type validation and characterisation of *An. gambaie* hemocyte lineages. **a**, UMAP visualisation of marker gene expression. Cells with more than 1 UMI shown in red. **b**, RNA *in-situ* hybridization combined with morphology analysis of circulating hemocytes. Actin shown in green, LRR8 in magenta, PPO4 in yellow, TM7318 in red and nuclei in blue. Scale bar: 5 µm. **c**, UMAP of 5383 hemolymph cells colored by cluster (cell state) identity with Seurat clustering. **d**, Unsupervised partition-based graph abstraction (PAGA) network analysis connecting stylised cell states based on UMAP clustering. Oen, oenocytoids; Div Gran, dividing granulocytes; Gran, granulocytes; Megac, megacytes; AM Gran, antimicrobial granulocytes; PHem, prohemocytes.

HC4 shares several markers with HC3, but is characterized by its own unique subset of markers (Fig. 1c). HC4 cells express cyclin B, aurora kinase and other mitotic markers, suggesting that they are proliferating hemocytes. Cells in HC5 and HC6 both are negative for PPO4 (Figs. 1c, 2a). HC5 cells express high levels of an uncharacterized transmembrane protein AGAP007318 (TM7318) and LPS-induced TNF-alpha transcription factor 3 (LL3) (Figs. 1c, 2a), while HC6 is negative for those markers and expresses antimicrobial peptides such as defensin 1 and cecropins 1 and C-type lysozyme (Figs. 1c). Cells in HC5 and HC6 are both LLR8^low^ and PPO4-negative but have two distinct morphologies (Fig. 2a-b). Cells negative for TM7318 (HC6) are small granulocytes that express antimicrobial genes (AM Gran) (16.4% of granulocytes), while TM7318 positive cells (HC5) are in low abundance (0.5% of granulocytes) and represent a new giant cell type (25-40 μm) that we named “megacytes” (Fig. 2b).

### Granulocyte lineages in *A. gambiae*

To investigate the differentiation dynamics of *A. gambiae* hemocytes, we re-clustered the cellular transcriptomes at higher resolution and performed lineage-tree reconstruction with partition-based graph abstraction (PAGA)(*16*). The major granulocyte population sub-clustered into three cell populations representing a basal, homeostatic state (Gran1), two states activated by blood-feeding and *Plasmodium*-infection (Gran2 and Gran3), (Fig. 2b-c). Prohemocytes were subclustered into two populations (PHem1 and PHem2), of which PHem2 appeared to be an intermediate stage between PHem1 and Gran1. We traced a connection between dividing granulocytes (Div Gran) and Gran3 granulocytes, which in turn link to Gran1. Gran1 also links to Gran2, which in turn links to megacytes (Megac), while Gran1 links directly to antimicrobial granulocytes (AM Gran; Fig. 2b-c). These findings were confirmed with two other cell-trajectory analysis packages: diffusion maps and slingshot (Fig. S4 and S5). Together, these analyses suggest the existence of a proliferative, oligopotent cell population that can replenish the pool of granulocytes and differentiate into more specialized hemocytes, such as megacytes and antimicrobial granulocytes. Granulocytes (HC4) express mitotic markers largely in response to blood feeding, in agreement with a recent report that blood feeding induced DNA synthesis in hemocytes(*17*). Our data suggests that granulocyte proliferation and prohemocyte differentiation both contribute to the observed increase in granulocyte numbers after blood feeding(*11*, *18*, *19*). However, the placement of prohemocytes in the granulocyte lineage tree should be treated with caution due to the paucity of unique markers.

Prohemocytes are proposed to be stem cells precursors of both granulocytes and oenocytoids(*11*). However, oenocytoids are transcriptionally disconnected from other hemocyte subtypes, and we did not observe transcriptional markers of cell proliferation in oenocytoids, suggesting they may represent a separate lineage which has its origin either in larval stages or in other adult tissues. Alternatively, oenocytoids could derive from granulocytes (Fig. 1b), but the differentiation rate may be very low when melanization responses are not elicited, which could result in few cells at intermediate stages of differentiation which may not be captured in our transcriptomic analysis.

To assess which of the newly discovered putative cell types are shared between anopheline and culicine mosquitoes, we also analyzed the single-cell transcriptome of 3123 cells from *A. aegypti*, a vector for several viral diseases including yellow fever, dengue, chikungunya and Zika. As with *Anopheles*, a dimensional reduction plot shows both canonical hemocytes and other cell types with mostly fat body signatures (Fig. 3a and Fig S6). A cross-species correlation analysis reveals two clusters (AaHC1 and AaHC2) with conserved transcriptome signatures for oenocytoids (99% and 77% correlation, respectively, with AgHC1) (Fig. 3a-b) and different granulocyte types, including antimicrobial peptide expressing cells (94% with AgHC6), and proliferating granulocytes (87% with AgHC4) (Fig. 3a-b; Table S4). Granulocytes and prohemocytes are again positioned on a continuum of transcriptomic similarity, with four different cell states, including a proliferating S-phase granulocyte cluster (AaHC6) without a clear *Anopheles* equivalent (Fig. 3a-b). Granulocytes express laminins, leucine-rich repeat proteins, scavenger receptors, Toll receptor 5, and the transcription factor Rel2 (Fig. S7 and Table S5). However, megacytes (AgHC5) lack an obvious counterpart in *Aedes*, and their unique marker (TM7318) is only present in anophelines of the *Cellia* subgenus (malaria vectors in Africa and Asia, Fig. S8).

**Fig. 3:**
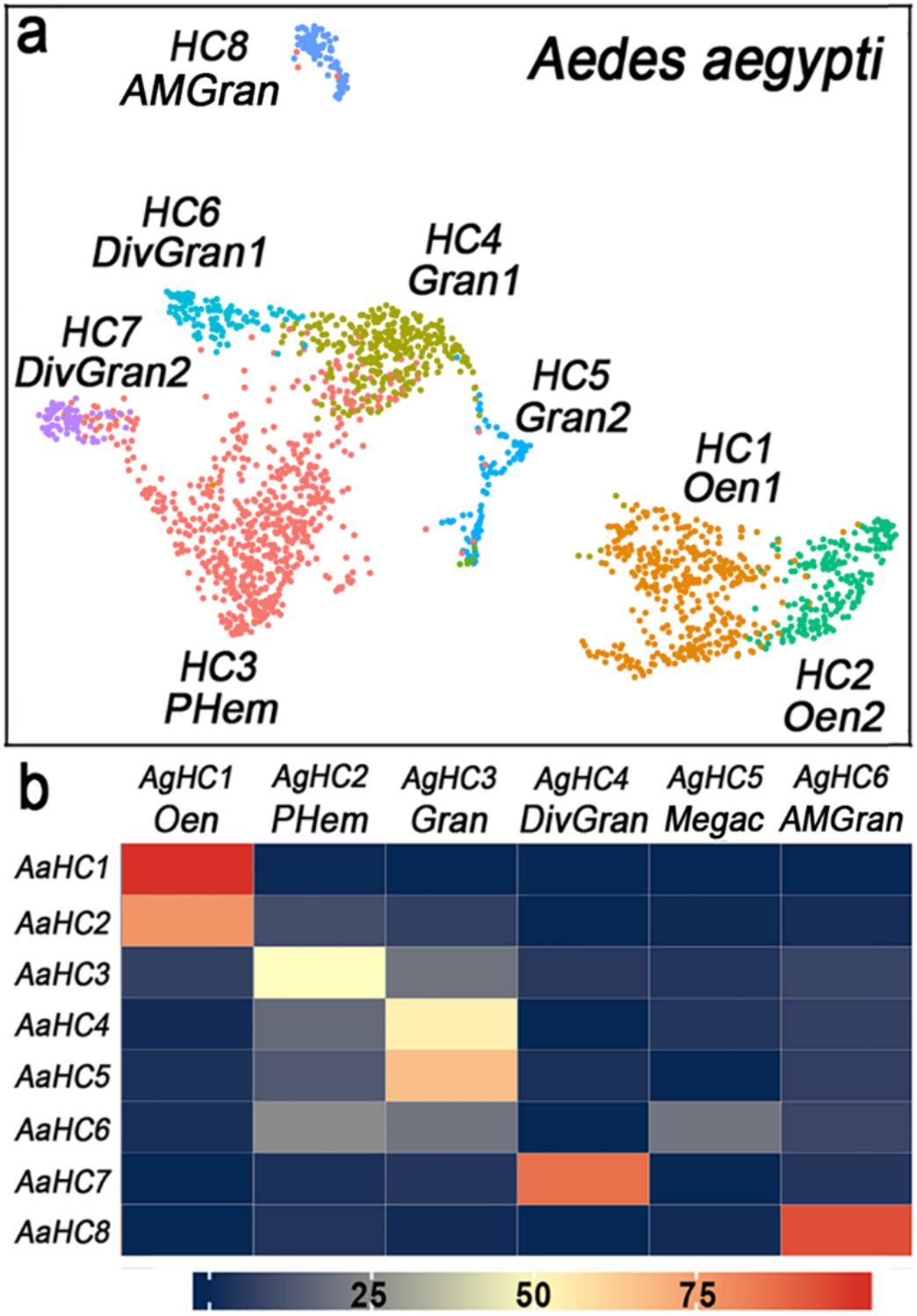
Characterisation *Aedes aegypti* hemocytes and correlation with *Anopheles* cell types. **a**, UMAP of 3123 *A. aegypti* hemocyte clusters colored by cluster identity with Seurat clustering. **b**, Heatmap showing probability of each *A. aegypti* hemocyte cell in the cluster belonging to each one of the *A. gambiae* (M-form) cell types after logistic regression and multinomial learning approach. *Ag*, *A. gambiae* (M-form); Aa, *A. aegypti*. Oen, oenocytoids; Div Gran, dividing granulocytes; Gran, granulocytes; Mega, megacytes; AM Gran, antimicrobial granulocytes; PHem, prohemocytes.

### Transcription factor LL3 is required for hemocyte differentiation during immune priming

The transcription factor LL3 can be detected in granulocytes from *Plasmodium*-infected *A. gambiae*, and silencing LL3 expression disrupts priming (*20*). However, it is not clear whether LL3 is essential for HDF synthesis or for hemocytes to respond to HDF. We found that LL3 is highly expressed in megacytes (HC5) (Figs. 1c and 4a) and explored whether silencing LL3 affects the HDF response. Transfer of hemolymph from primed *A. gambiae* donors had HDF activity and elicited a strong priming response in control recipients injected with *lacZ* double stranded (ds) RNA, resulting in a prominent increase in circulating granulocytes, a modest increase in oenocytoids and a decrease in prohemocytes (Fig. 4b and Table S6-9). This response was completely abolished when LL3 expression was silenced in the recipients by injection of dsLL3 RNA (Fig. 4b and Table S6-9), indicating that LL3, and probably megacytes, play a key role in orchestrating hemocyte responses to HDF.

**Fig. 4:**
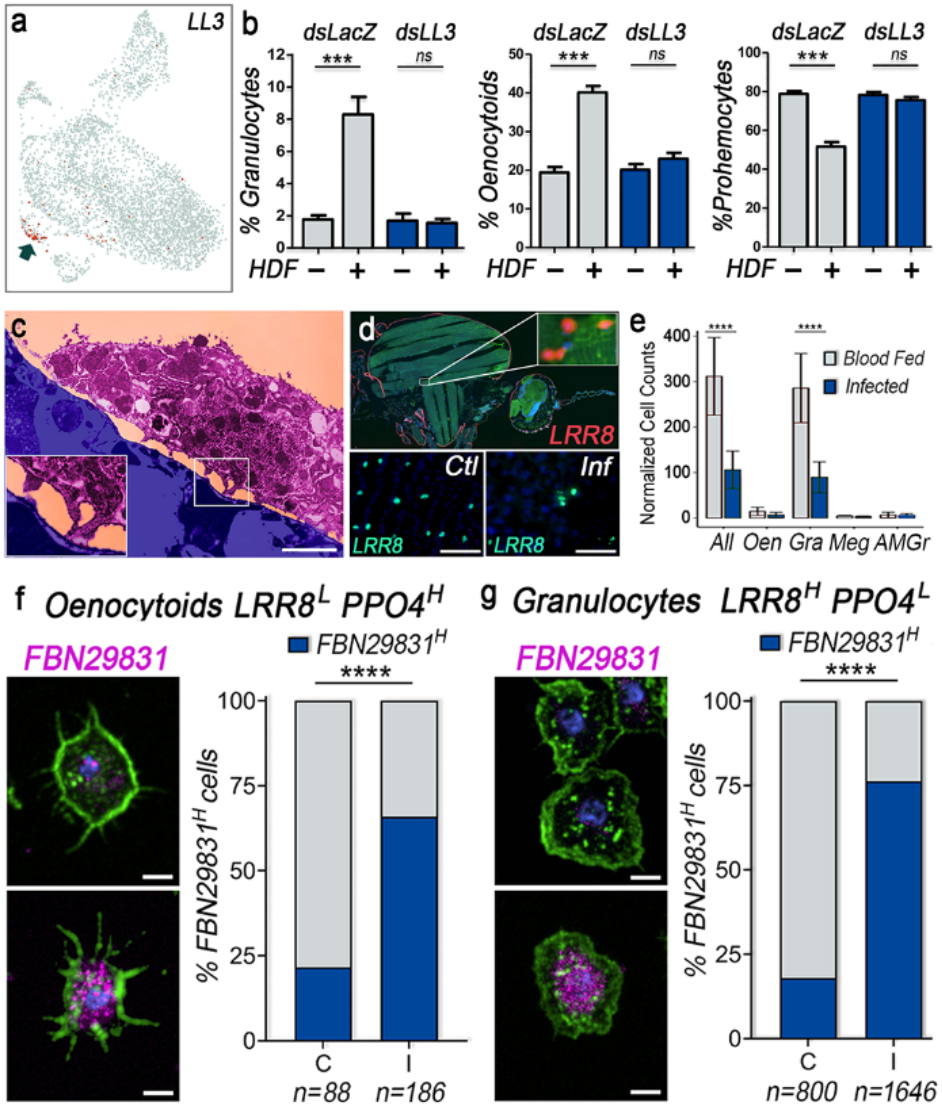
*Anopheles gambiae* cellular immune responses to *Plasmodium* infection. **a**, UMAP visualisation of all hemocytes by *LL3* expression. Red indicates cells with more than 1 UMI. **b**, Percentage of circulating granulocytes, oenocytoids and prohemocytes of *LL3*-silenced mosquitoes injected (+) or not (−) with HDF versus double-stranded *lacZ* RNA injected mosquitoes used as negative controls (**** P<0.0001, Unpaired t-test). Data are representative of two independent experiments (mean ± SEM). **c**, Transmission electron microscopy (TEM) in false colors depicting *A. gambiae* (M form) granulocyte (purple) attached via pseudopodia (insert) to the abdominal fat body (blue); scale bar: 1.5 µm. **d**, RNA *in-situ* hybridisation of a longitudinal section (top panel) or carcass whole-mounts (bottom panels) of control blood-feed (Ctl) and *P. berghei* (Inf) infected *A. gambiae* mosquitoes. Top panel: GAPDH shown in green, LRR8 in red. Bottom panels: LRR8 in green. Scale bar: 20 µm. **e**, Quantification of *Anopheles* hemocytes attached to the mosquito fat body with blood-feeding or *P. berghei* infection, normalised by the surface area of the fat bodies. RNA *in-situ* hybridisation of a longitudinal section (top panel) or carcass whole-mounts. All, all hemocytes; Oen, oenocytoids; Gran, granulocytes; AM Gran, antimicrobial granulocytes. Data are representative of three independent experiments merged (mean ± SEM, **** P<0.0001, Welch T-test). **f**, Percentage of circulating oenocytoids (LRR^L^PPO4^H^) and **g**, granulocytes (LRR^H^PPO4^L^) positive for *FBN29831* in control (C) and *P. berghei* infected (I) mosquitoes, 48 hours post feeding (right panels). Representative RNA *in-situ* hybridisation pictures of oenocytoids (**f**) and granulocytes (**g**) with low and high expression of FBN29831 (left panels). Actin is shown in green, FBN29831 in magenta, nuclei in blue. Scale bar: 5 µm. Data are representative of two independent experiments merged (****P<0.0001, Chi-square test).

### Blood-feeding and *Plasmodium* infection trigger granulocyte activation and mobilization

The proportion of circulating granulocytes has been shown to be low (1-3%) under normal conditions but increases in response to *Plasmodium* infection(*2*). Whether the increase is due to granulocyte proliferation, differentiation from prohemocytes or to mobilization of sessile hemocytes was unknown. Transmission electron microscopy of sugar fed mosquitoes showed individual sessile hemocytes bathed by hemolymph and attached to the basal lamina of the tissues through pseudopods (Fig. 4c), indicative of a dynamic and potentially transient association. Using whole tissue mount *in situ* hybridization we found that most sessile hemocytes are PPO4^low^/LLR8^high^ granulocytes (89.3 ± 6.2%), while PPO4^high^/LLR8^low^ oenocytoids are less abundant (4.2 ± 3.1%) and TM7318 positive megacytes are even rarer (2.7 ± 2.3%) (Fig. 4d-e and Table S10-11). Furthermore, we found a dramatic reduction of sessile PPO4^low^/LLR8^high^ granulocytes in response to *Plasmodium* infection (P<0.0001, Welch T-Test, Fig. 4e and Table S12-13), while there was no significant difference in the numbers of sessile PPO4^high^/LLR8^low^ oenocytoids, TM7318^+^ megacytes or TM7318^−^ AM granulocytes (Fig. 4e and Table S12-13). In circulating hemocytes, *P. berghei* infection induced high expression of a fibrinogen-like protein AGAP029831 (FBN29831) (Fig. S9). RNA *in situ* hybridization showed that the proportion of FBN29831-positive cells increased in both PPO^low^/LLR8^high^ granulocytes (P<0.0001, X^2^, Fig. 4f and Table S14-15) and PPO4^high^/LLR8^low^ oenocytoids (P<0.0001, X^2^, Fig. 4g and Table S16-17), indicating that this is a general marker of hemocyte immune activation. Infecting *A. gambiae* with the human parasite *P. falciparum* produced a similar increase in FBN29831-positive cells (Fig. S10). Combined, our results suggest that hemocyte recruitment from the body wall, granulocyte activation and proliferation, and prohemocyte differentiation can all contribute to boost circulating granulocyte numbers upon immune challenge.

Our knowledge of cellular immunity in vertebrates relies critically on understanding the functional diversity of cell types, their developmental trajectories and their trafficking dynamics. The hemocyte atlas presented in this study represents significant progress towards this level of understanding for two invertebrate immune systems that limit the vectorial capacity of mosquitoes for a range of important human pathogens. We confirm the existence of two major canonical hemocyte types in mosquitoes, the oenocytoids and granulocytes, and with the help of transcript markers we relate cellular morphology to a more comprehensive molecular characterization of these cells. Unlike current thinking in the field we show prohemocytes and granulocytes to be closely related cells, and identified transcriptional profiles and molecular markers that define previously unknown hemocyte subtypes (megacytes, proliferating granulocytes, and antimicrobial granulocytes), as well as a subpopulation of fat body cells (AgFBC1) with potential immune-modulatory functions.

We define two main hemocyte lineages in *A. gambiae*: the oenocytoid lineage, and the prohemocyte-granulocyte lineage. The latter can be further split into three sub lineages leading to differentiated megacytes, antimicrobial granulocytes, or proliferating granulocytes. We discovered well-defined molecular diversity within these cell types and show them to be largely conserved between distantly related mosquito genera and presumably of functional importance. Conversely, megacytes were not detected in our *A. aegypti* mosquito dataset. Megacytes have a unique morphology, and silencing of transcription factor LL3 provides tentative evidence for a regulatory role of these cells in HDF mediated immune priming. However, in another study LL3 was found upregulated more broadly in granulocytes after *Plasmodium* infection(*20*), and we cannot rule out with certainty that other hemocyte types express LL3 at levels below the detection limit of scRNA-seq, and might thus be directly affected by LL3 silencing.

The cell-type markers and FISH probes we identified and validated will allow investigators to probe the immune functions of megacytes and other specialized hemocyte types in detail. Our analysis cannot resolve the developmental origin of oenocytoids, but we identify two potential origins for the observed expansion of circulating granulocytes upon blood feeding. One is the mobilization of sessile PPO4^low^/LLR8^high^ cells from the body wall, the other is a pool of proliferating, oligopotent granulocytes. Whether prohemocytes, which appear transcriptionally related to but less responsive than granulocytes, can transform into granulocytes and whether they need to enter the cell cycle, remains to be determined.

In sum, the cell-type-specific marker genes, reference transcriptomes, and companion website (https://hemocytes.cellgeni.sanger.ac.uk/) from our study will serve as a resource for the field, and provide a starting point for the type of lineage tracing and functional experiments which, in vertebrates, are resolving the developmental origins and functions of diverse immune cell populations.

## Acknowledgements

The authors are grateful to Tom Metcalf for technical support, to Andrew Goldsborough for making available the vivoPHIX reagent and advising on its use, to Jana Eliasova for help with illustrations. This work was supported by the Intramural Research Program of the Division of Intramural Research Z01AI000947, National Institute of Allergy and Infectious Diseases (NIAID), NIH to CBM, by an NIH Oxford-Cambridge scholarship, the UCLA-Caltech MSTP, and the NIGMS T32 GM008042 to GR, Wellcome core grant 206194/Z/17/Z to the Sanger Institute and funding from the Knut and Alice Wallenberg foundation and the European Research Council (Grant agreement No. 788516) to OB.

## Ethics Statement

Public Health Service Animal Welfare Assurance #A4149-01 guidelines were followed according to the National Institutes of Health Animal (NIH) Office of Animal Care and Use (OACU). These studies were done according to the NIH animal study protocol (ASP) approved by the NIH Animal Care and User Committee (ACUC), with approval ID ASP-LMVR5.

## References

1. B. C. Ho, E. H. Yap, M. Singh, Melanization and encapsulation in Aedes aegypti and Aedes togoi in response to parasitization by a filarioid nematode (Breinlia booliati). Parasitology. 85 (Pt 3), 567–575 (1982).

2. J. Rodrigues, F. A. Brayner, L. C. Alves, R. Dixit, C. Barillas-Mury, Hemocyte differentiation mediates innate immune memory in Anopheles gambiae mosquitoes. Science. 329, 1353–1355 (2010).

3. WHO, World malaria report 2019 (https://www.who.int/publications-detail/world-malaria-report-2019).

4. S. Luckhart, Y. Vodovotz, L. Cui, R. Rosenberg, The mosquito Anopheles stephensi limits malaria parasite development with inducible synthesis of nitric oxide. Proc. Natl. Acad. Sci. U. S. A. 95, 5700–5705 (1998).

5. S. Blandin, S.-H. Shiao, L. F. Moita, C. J. Janse, A. P. Waters, F. C. Kafatos, E. A. Levashina, Complement-like protein TEP1 is a determinant of vectorial capacity in the malaria vector Anopheles gambiae. Cell. 116, 661–670 (2004).

6. J. L. Ramirez, L. S. Garver, F. A. Brayner, L. C. Alves, J. Rodrigues, A. Molina-Cruz, C. Barillas-Mury, The role of hemocytes in Anopheles gambiae antiplasmodial immunity. J. Innate Immun. 6, 119–128 (2014).

7. J. C. Castillo, A. B. B. Ferreira, N. Trisnadi, C. Barillas-Mury, Activation of mosquito complement antiplasmodial response requires cellular immunity. Sci. Immunol. 2 (2017), doi:10.1126/sciimmunol.aal1505.

8. S. Kumar, L. Gupta, Y. S. Han, C. Barillas-Mury, Inducible peroxidases mediate nitration of anopheles midgut cells undergoing apoptosis in response to Plasmodium invasion. J. Biol. Chem. 279, 53475–53482 (2004).

9. G. de A. Oliveira, J. Lieberman, C. Barillas-Mury, Epithelial nitration by a peroxidase/NOX5 system mediates mosquito antiplasmodial immunity. Science. 335, 856–859 (2012).

10. J. L. Ramirez, G. de Almeida Oliveira, E. Calvo, J. Dalli, R. A. Colas, C. N. Serhan, J. M. Ribeiro, C. Barillas-Mury, A mosquito lipoxin/lipocalin complex mediates innate immune priming in Anopheles gambiae. Nat. Commun. 6, 7403 (2015).

11. J. C. Castillo, A. E. Robertson, M. R. Strand, Characterization of hemocytes from the mosquitoes Anopheles gambiae and Aedes aegypti. Insect Biochem. Mol. Biol. 36, 891–903 (2006).

12. J. G. King, J. F. Hillyer, Infection-induced interaction between the mosquito circulatory and immune systems. PLoS Pathog. 8, e1003058 (2012).

13. J. G. King, J. F. Hillyer, Spatial and temporal in vivo analysis of circulating and sessile immune cells in mosquitoes: hemocyte mitosis following infection. BMC Biol. 11, 55 (2013).

14. G. M. Attardo, I. A. Hansen, A. S. Raikhel, Nutritional regulation of vitellogenesis in mosquitoes: implications for anautogeny. Insect Biochem. Mol. Biol. 35, 661–675 (2005).

15. M. S. Severo, J. J. M. Landry, R. L. Lindquist, C. Goosmann, V. Brinkmann, P. Collier, A. E. Hauser, V. Benes, J. Henriksson, S. A. Teichmann, E. A. Levashina, Unbiased classification of mosquito blood cells by single-cell genomics and high-content imaging. Proc. Natl. Acad. Sci. U. S. A. 115, E7568–E7577 (2018).

16. F. A. Wolf, F. K. Hamey, M. Plass, J. Solana, J. S. Dahlin, B. Göttgens, N. Rajewsky, L. Simon, F. J. Theis, PAGA: graph abstraction reconciles clustering with trajectory inference through a topology preserving map of single cells. Genome Biol. 20, 59 (2019).

17. W. B. Bryant, K. Michel, Blood feeding induces hemocyte proliferation and activation in the African malaria mosquito, Anopheles gambiae Giles. J. Exp. Biol. 217, 1238–1245 (2014).

18. L. A. Baton, A. Robertson, E. Warr, M. R. Strand, G. Dimopoulos, Genome-wide transcriptomic profiling of Anopheles gambiae hemocytes reveals pathogen-specific signatures upon bacterial challenge and Plasmodium berghei infection. BMC Genomics. 10, 257 (2009).

19. J. Castillo, M. R. Brown, M. R. Strand, Blood feeding and insulin-like peptide 3 stimulate proliferation of hemocytes in the mosquito Aedes aegypti. PLoS Pathog. 7, e1002274 (2011).

20. R. C. Smith, C. Barillas-Mury, M. Jacobs-Lorena, Hemocyte differentiation mediates the mosquito late-phase immune response against Plasmodium in Anopheles gambiae. Proc. Natl. Acad. Sci. U. S. A. 112, E3412–3420 (2015).

