## Supplementary material for "Mosquito cellular immunity at single-cell resolution": Methods and Supplementary Figures and Tables

**Figure S1. Quality control (QC) of *Anopheles gambiae* hemocyte scRNA-seq data.**
**a**, Cell numbers after QC by condition. **b**, UMAP dimensionality reduction plot coloured by experiments and samples showing mixing after data integration with Seurat 3.0. **c**, QC metrics for the integrated dataset. Genes = number of genes detected per cell. Mt = proportion of reads mapping to mitochondrial genes.

**a**

| *Days* | 0 | 1 | 2 | 3 | 7 |
|---|---|---|---|---|---|
| **Sugar** | 1049 | n/a | n/a | n/a | n/a |
| **Blood** | n/a | 78 | 707 | 661 | 257 |
| **P. berghei** | n/a | 425 | 270 | 911 | 1025 |

**b**

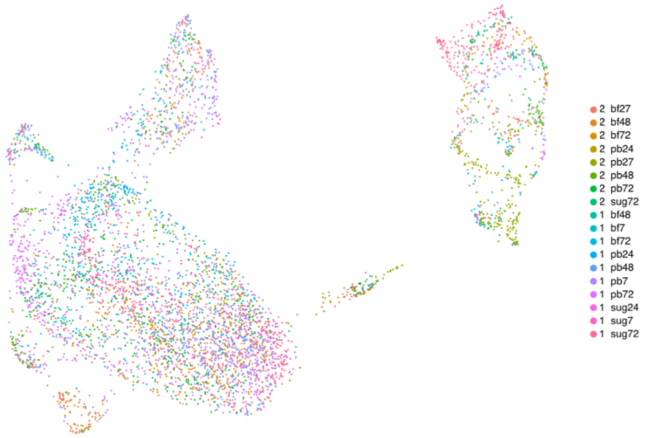


**c**

**d**

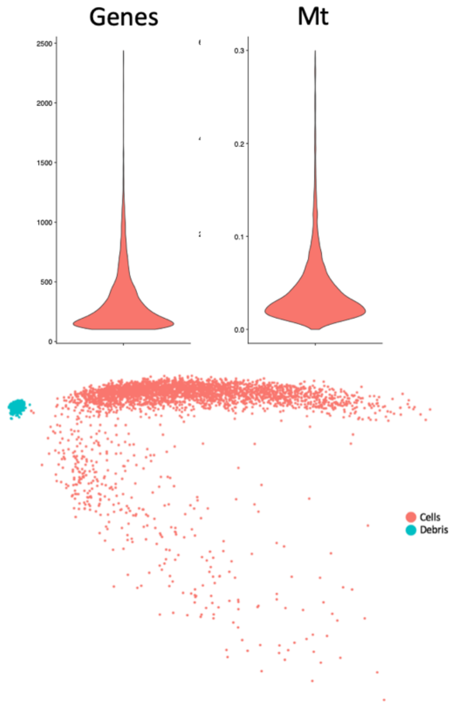

**Figure S2. Heatmap of top ten gene markers for each *A. gambiae* cell cluster.** DE genes were identified with the Wilcoxon rank-sum test. P-values were adjusted for multiple testing using the Bonferroni correction. All P-adjusted values < 0.001, with genes ordered by average log fold change between cluster of interest and all other cells. Down-sampled to 300 cells per cluster for clarity. Scale from yellow (high expression) to pink (low expression).

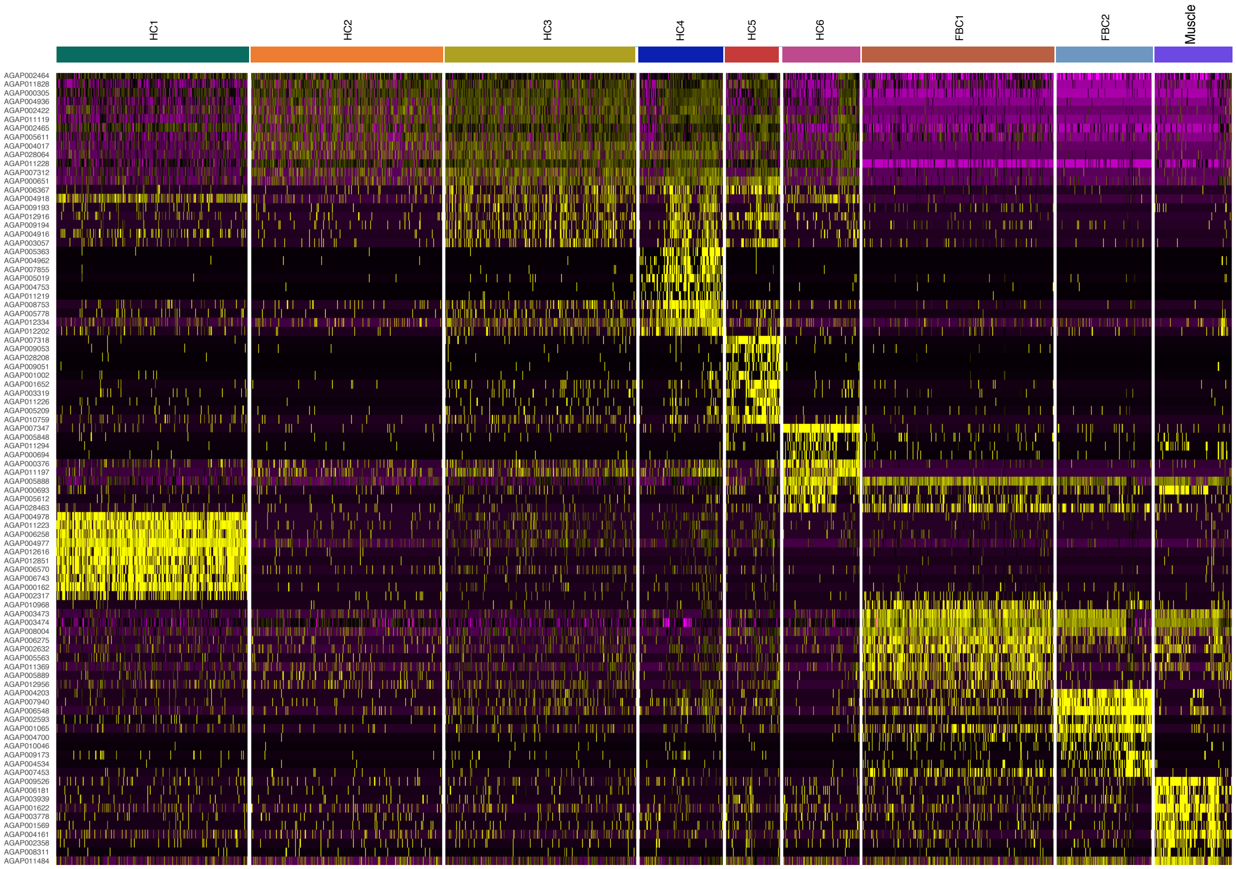

**Figure S3.** ***A. gambiae*** **bulk RNAseq data. a**, Differential expression analysis of hemocytes vs average of mosquito guts and carcasses with DEseq2, filtered for $log_2$-fold change > 2 and Wald significance testing Q < 0.001. Top 5 marker genes of each scRNA-seq clusters are displayed. **b**, Distance matrix correlating the overall similarity and hierarchical clustering of each sample. **c**, PCA analysis and clustering of samples based on overall transcriptional similarity, divided by sample type and experiment. Carc = carcasses; Gut = guts; Hem = hemocytes. Data from three independent biological replicates for each condition and time-point (day 0, 1, 2, 3, and 7 after sugar-feeding, blood-feeding or *P. berghei* infection).

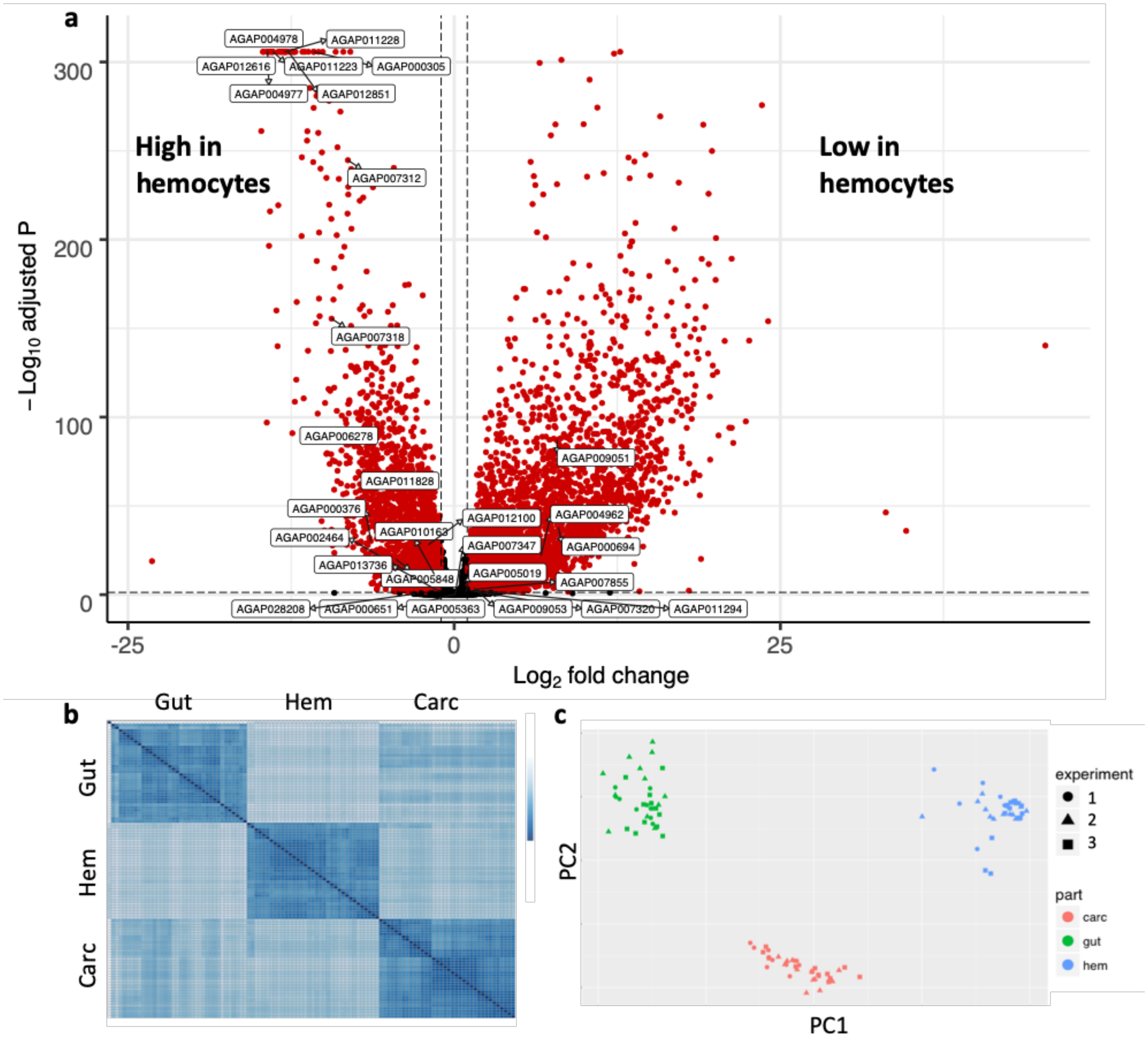

**Figure S4. Hemocyte lineage tracing.**

**a,** 2D diffusion map of granulocyte lineages. **b,** 3D diffusion map of granulocyte lineage. **c,** DC1 vs. DC3 plot highlights transition between Div Gran and Gran3. **d,** DC1 vs. DC3 plot highlights transition between Megac and Gran2. **e,** DC2 showcases hemocyte maturity, with proliferating cells on the right and differentiated states on the left.

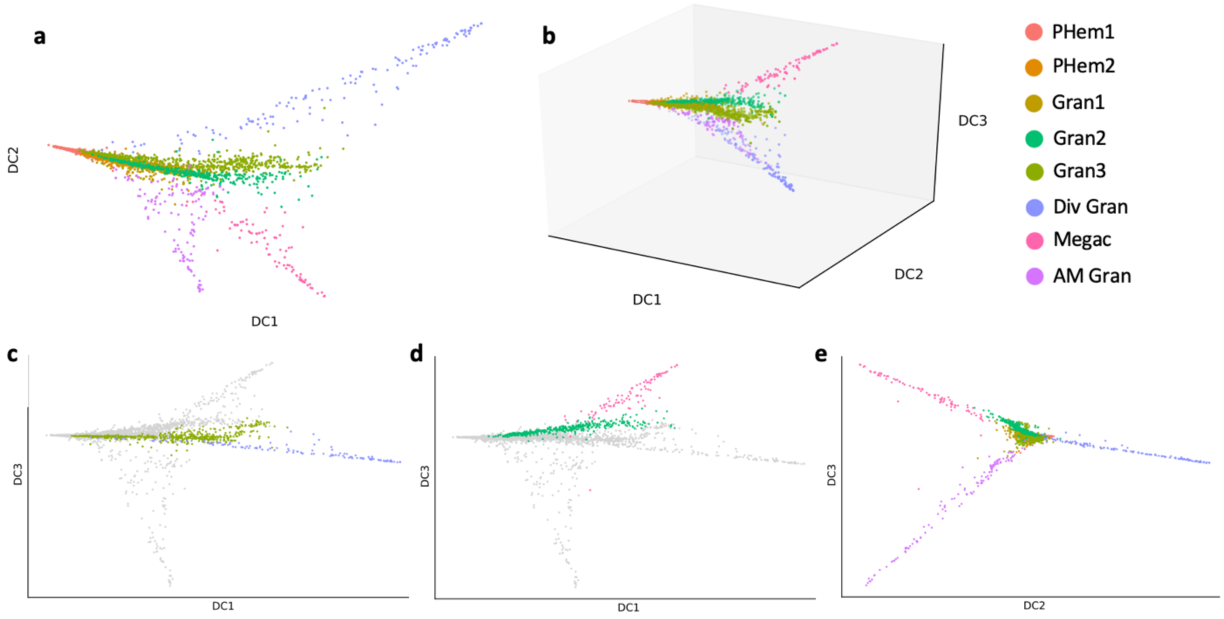

**Figure S5. Lineage tracing and pseudotime reconstruction of granulocytes and prohemocytes.**
**a,** Slingshot analysis after subsetting non-hemocytes and oenocytoids. **b,** Pseudotime reconstruction on DC1 vs. DC2. **c-e,** Pseudotime reconstruction with Slingshot for each separate lineages from ProH to **(c)** Div Gran **(d)** Megac **(e)** AM Gran.

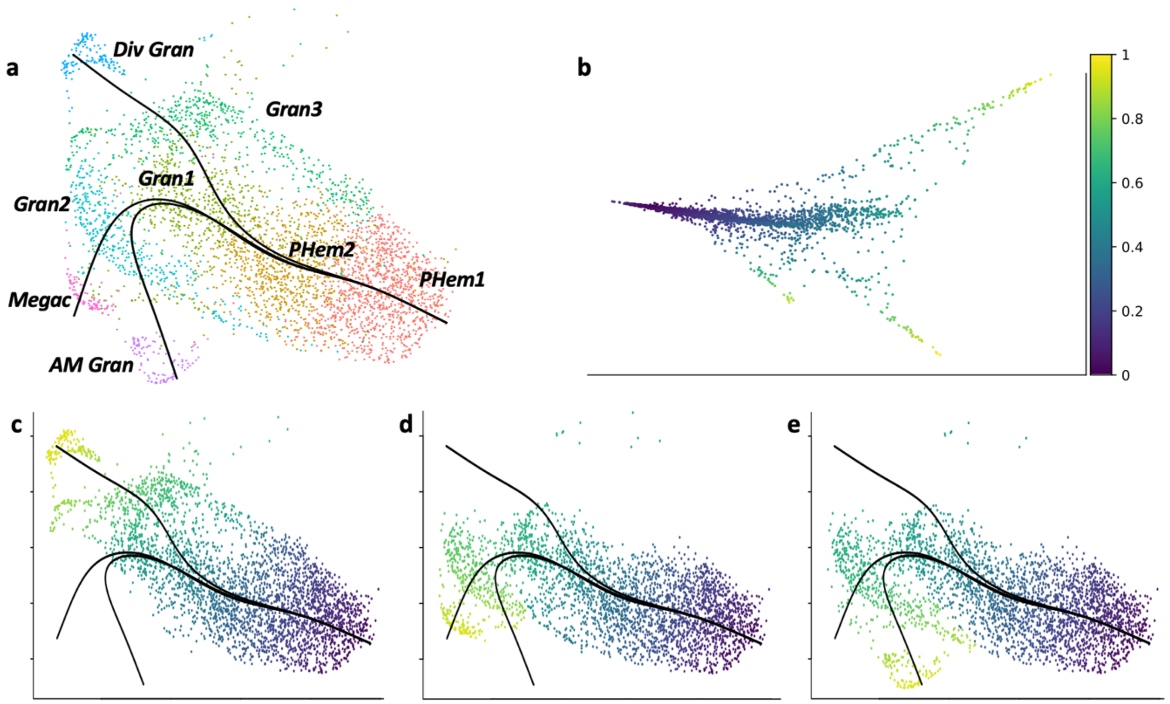

**Figure S6. Quality control (QC) of *A. aegypti* hemocyte scRNA-seq data**

**a**, Full UMAP dimensionality reduction plot coloured by cell cluster. **b**, UMAP dimensionality reduction plot coloured by experiments and samples showing mixing after data integration with Seurat 3.0. **c**, table of cells after QC per condition. **d**, QC metrics for the integrated dataset. Genes = number of genes detected per cell. Mt = proportion of reads mapping to mitochondrial genes. **e**, *A. aegypti* hemocyte morphology. Stains stained with phalloidin (actin) in green and Hoechst (nuclei) in blue. Scale bar: 5 µm.

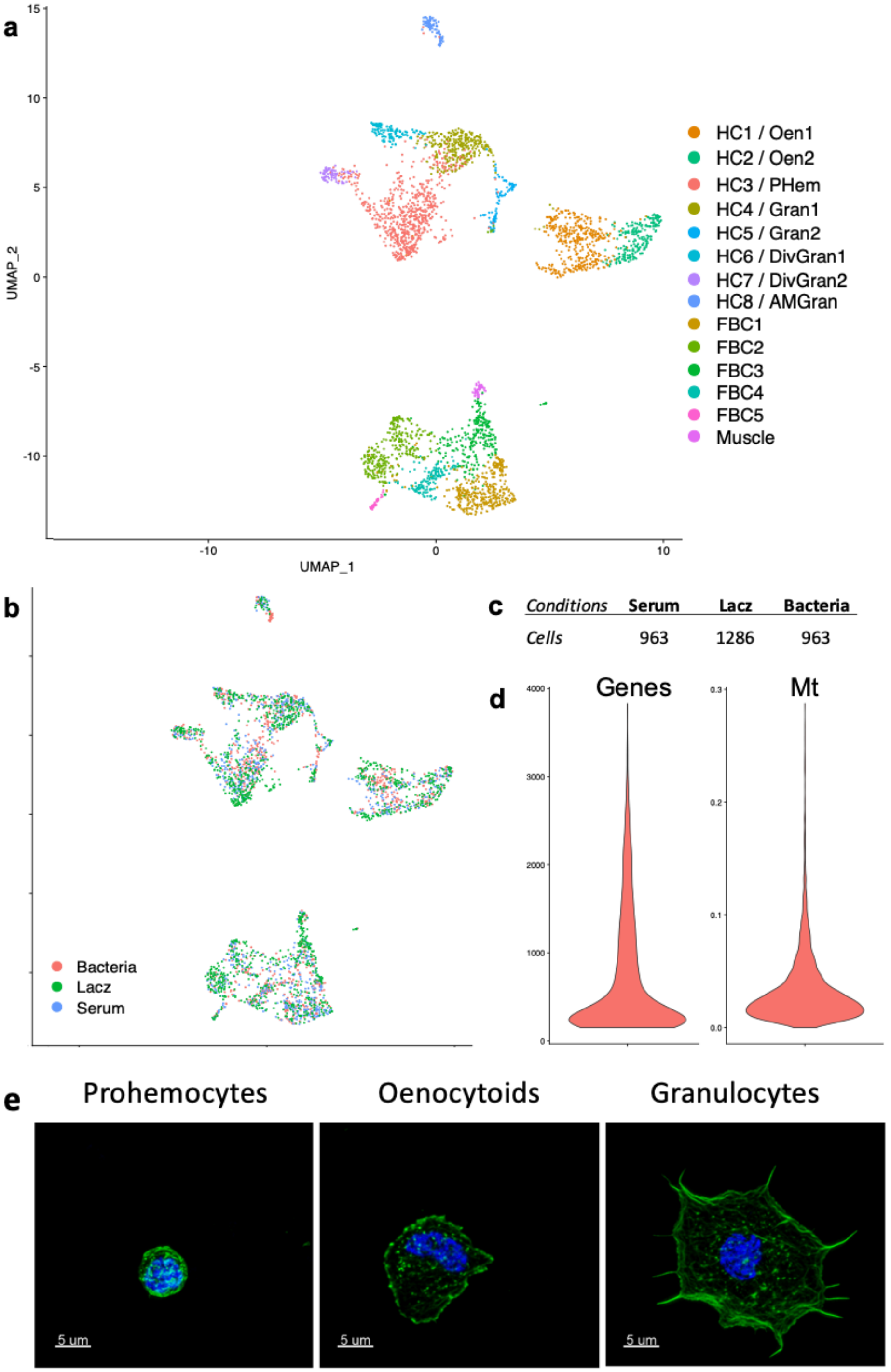

**Figure S7. Heatmap of top ten gene markers of each *A. Aegypti* cell cluster**

DE genes were identified with the Wilcoxon rank-sum test. P-values were adjusted for multiple testing using the Bonferroni correction. All P-adjusted values < 0.001, with genes ordered by average log fold change between cluster of interest and all other cells. Down-sampled to 300 cells per cluster for clarity. Scale from yellow (high expression) to pink (low expression).

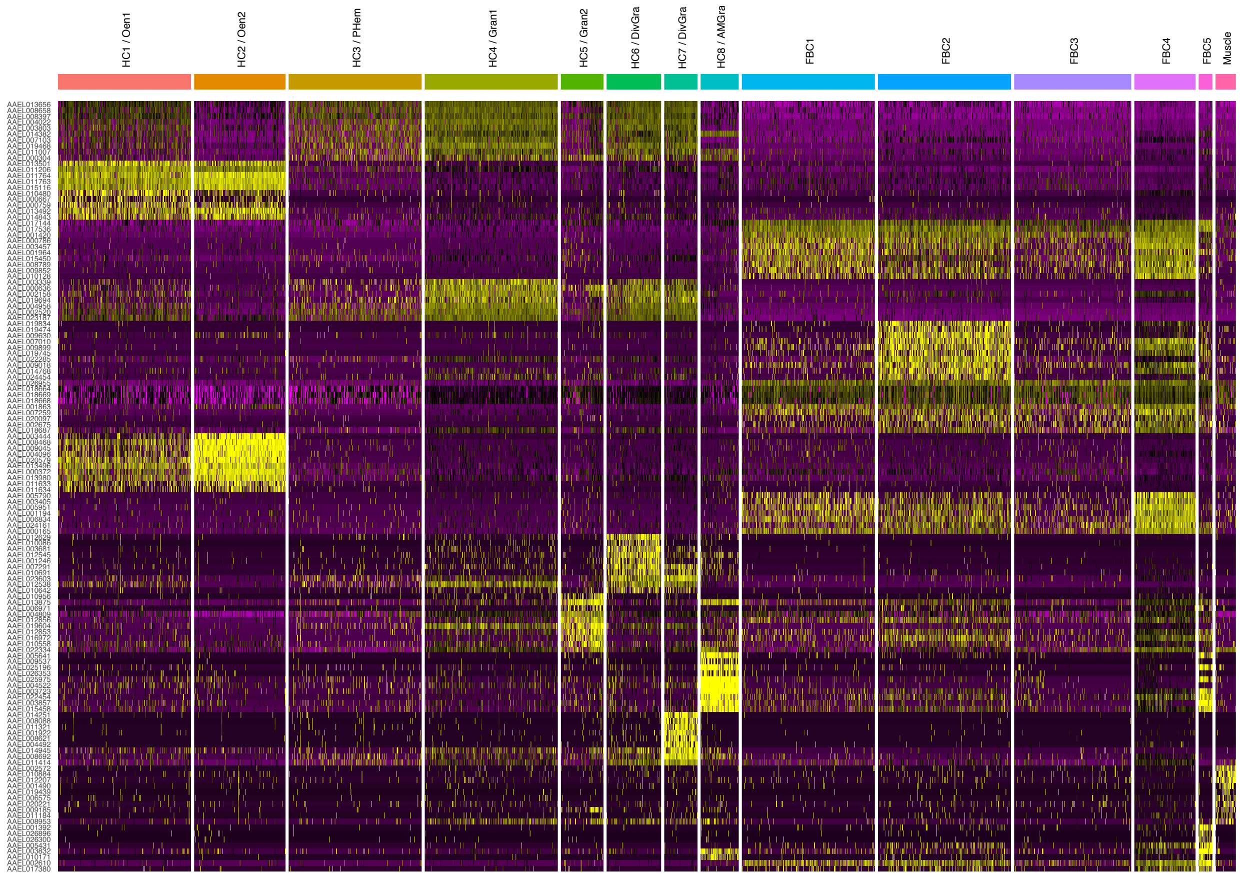

**Figure S8. Phylogeny of the megacyte marker, TM7318, limited to the *Cellia* subgenus.** Multiple sequence alignment (A) and neighbor-joining phylogenetic tree (B) (without distance correction) of the amino acid sequence the *Anopheles gambiae* protein (AGAP007318) and putative orthologs from other anopheline mosquitoes of the subgenus *Cellia. (Amel) A. melas; (Agam) A. gambiae; (Aara) A. arabiensis; (Aepi) A. epiroticus; (Adi), A.dirus; (Aste) A. stephensi; (Afar) A. farauti.*

**A)** CLUSTAL Omega (1.2.4) multiple sequence alignment

```
Agam      MGLPTFLNFNKNIQKHAMTVGVILTLYTIFTLLIGVAWLIEIKEQASLYKLDFRVDLTGI    60
Amel      MGLPTFLSFNKNIQKHALSVGVILTLYTIFTLLIGVAWLIEIKEQVSLYKLDFRVDLTGI    60
Aara      MGLPTFLNFNKNIQKHAMAVGIILTLYTIFTLLIGVAWLIEIKEQATLYKLDFRVDLTGI    60
Aepi      MGLPAFLDFSSKIRLHATIVGMLFTLDTIVTLALGIALPVEGKHVPPEL-----------    49
Adir      MKLPSFLDFSANIRTHGAFVGVLLIAITIPTLVLAAVSNFRTEDLPVEV-----------    49
Aste      MALPKCLNFSNNIRTHAMFVGVIVSLDAICILLFSAAWNIKAEQLPKEL-----------    49
Afar      MALPSYLDFSANIRTHAAFVGVLLALDAIITLLLSAVWNIKAENLPDEF-----------    49
          * **  *.*. :*: *.  **::.   :*  * :. .  .. :.

Agam      RTINPRMAGFWEKEGFLVMGIGFAILAVLYWITYLMRQRLLLAVFIGLMVIIVTLNVIGM    120
Amel      RTINPRMAGFWEKEGFLVMGIGFAILTVLYCVTYLMRQRLLLAVFIGLMVIIITLNIIGM    120
Aara      RTINPRTAGFWEKEGFLVMGIGFAILAVLYWITYLMRQRLLLAVFIGLMVIIVTLNIIGM    120
Aepi      ---APHM--HVIKNFLIVAGVLNALFTLMYWVGFLKRQRWCLTVCIGFEAVAITLQFISL    104
Adir      ----KHI--GYMHEAVLIAGTVKILFTVIFWIGFLKRKRWCLTVFVVFLAVALTLLYLGL    103
Aste      ----QHL--ASMREVLLVMGLINAFFALVYWVGFLKRQRWCLTLFIGFLAVAITLHFIGF    103
Afar      ----KHI--ANLRQVLLILGIGEALFATLYWVGFLKRVRWCLTIVVGFLAVVLTLQFIGL    103
               :      :: .:: *    ::: :: : :* * *  *:: : : .: :**  :.:

Agam      VAAGLKLKFVLACVVMFGLPVNVYMLVVALQLHKSKFIFTPRQQG    165
Amel      VAAGLKLKFVLACVVMFGLPVNVYMLLVARQVHKSKFTFTARQQG    165
Aara      VAAGLKLKFVLACVVMFGLPVNVYMLVVALQLHKSKFIFTTRQQG    165
Aepi      LGAMLQLNAVGFLGCLIALALNGYVLLVTLQLRKANSNLSVTQQG    149
Adir      LGAILLQHFLAVCFMLIALAASKYVLLVTLQLREVNSNEMVPLER    148
Aste      LGAILQLQIVSACAILIVLGINGYVLLVTLQLRKANLDSPLSQDG    148
Afar      LGALLQLNIASACMVLIVLAVNGYVLLVTLQLRKANSIVTVPQEA    148
          :.* *  :       :: *  . *:*:*: *::: :       :
```

**B)**

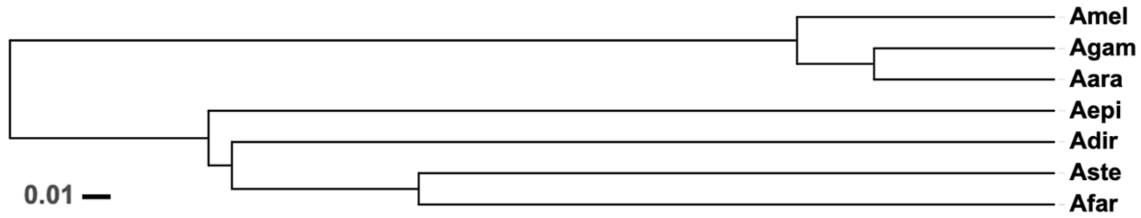

**Figure S9. FBN29831 gene expression in hemocytes in control blood-fed and *P. berghei* infected mosquitoes 48 hours post feeding.** Hemocytes from mosquitoes fed in an uninfected mouse (C) and in a *P. berghei* infected mouse (I). Experiment was performed with 3 biological replicates with 20 mosquitoes each were used per condition. Unpaired t-test was used for statistical analysis. *p < 0.0139

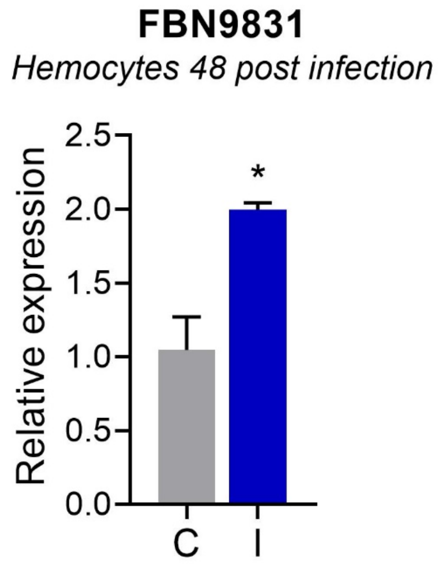

**Figure S10. FBN29831-positive cells in *A. gambiae* G3 after *Plasmodium falciparum* infection.** Percentage of circulating hemocytes positive for *FBN29831* in control blood fed (C) and *P. falciparum* infected (I) mosquitoes, 48 hours post feeding (left panels) as determined by microscopic analysis of circulating hemocytes from batches of 8-10 mosquitoes labelled by RNA-FISH. Table show percentages and numbers of hemocytes expressing high and low levels of FBN29831 in control and infected mosquitoes. Data are representative of 2 independent experiments. Chi square test was used for statistical analysis.

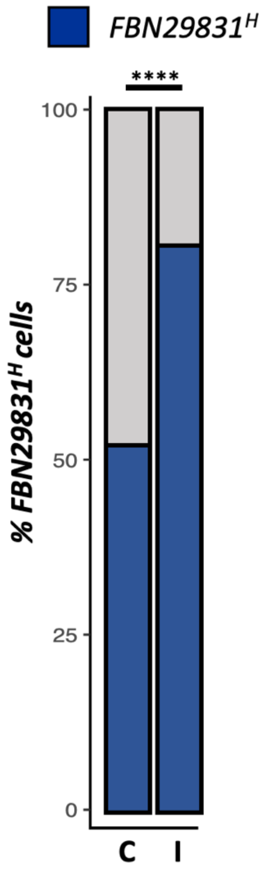


| | | Rep1 | | Rep2 | | Merge | |
|---|---|---|---|---|---|---|---|
| | | **%** | **# cells** | **%** | **# cells** | **%** | **# cells** |
| Control | FBN29831L | 50.2 | 271 | 55.5 | 292 | 52.8 | 563 |
| | FBN29831H | 49.8 | 269 | 44.5 | 234 | 47.2 | 503 |
| Infected | FBN29831L | 11.4 | 35 | 19.4 | 134 | 17.0 | 169 |
| | FBN29831H | 88.6 | 271 | 80.6 | 556 | 83.0 | 827 |

| | **C vs I - LRR FBN Adjusted Chi square** |
|---|---|
| Chi-square, df | 288.957, 1 |
| z | 17.00 |
| P value | <0.0001 |
| P value summary | **** |

### Supplementary Tables

**Table S1**. Marker genes of *Anopheles gambiae* cell clusters. P val adj = P-value adjusted for multiple testing. Avg $log_2$FC = average $log_2$ fold change between cluster of interest and all other clusters. Pct. 1 = percentage of cells in the cluster with detectable marker expression. Pct. 2 = percentage of cells in all other clusters with detectable marker expression. Electronic annotation.

(see separate file)

**Table S2**. Differentially expressed genes in hemocytes from bulk RNAseq analysis of *Anopheles gambiae* tissues. Differential expression analysis in hemocytes vs. the average expression of mosquito carcasses and guts was performed with DEseq2, genes filtered for $log_2$ fold change > 1 and Wald significance testing of Q < 0.05. Mean is the mean of normalized counts of all samples, normalized for sequencing depth. $Log_2$ FC (fold change) is the effect size estimate. On a logarithmic scale to base 2. Lfc SE is the standard error estimate for the $log_2$FC estimate. Data representative of 3 independent experiments

(see separate file)

**Table S3**. RNA-FISH markers chosen by virtue of their total expression and expression specificity in scRNA-seq and bulk RNAseq (+/++/+++ qualitative assessment).

| Markers | scRNA - specificity | scRNA - expression | Bulk vs gut - log2 fold | Bulk vs carcass - log2 fold | Description |
|---|---|---|---|---|---|
| ***General*** | | | | | |
| AGAP009623 | n/a | n/a | n/a | n/a | GAPDH - mosquito pos. control |
| AGAP008296 | n/a | n/a | -13.2 | -7.6 | Trypsin - gut |
| AGAP004203 | +++ | +++ | 4.1 | -2.5 | Vitellogenin - FBC2 |
| ***HC1*** | | | | | |
| AGAP012851 | +++ | +++ | 6.9 | 4.7 | PPO4 |
| AGAP012000 | ++ | ++ | 8.1 | 5.5 | Aldo-keto-reductase |
| ***HC2 and HC3*** | | | | | |
| AGAP004017 | ++ | +++ | 7.3 | 4.8 | LRR8 |
| AGAP011974 | ++ | ++ | 5.6 | 4.2 | SCRC1 |
| ***HC5*** | | | | | |
| AGAP007318 | +++ | ++ | 5.3 | 2.8 | TM7318 |

**Table S4**. Probability that each cell in a mosquito cluster belongs to another cell cluster in the opposite species (*Anopheles gambiae* vs *Aedes aegypti*) after logistic regression with L2-norm regularization and a multinomial learning approach on log-transformed normalized data.

| | *AaHC1* | *AaHC2* | *AaHC3* | *AaHC4* | *AaHC5* | *AaHC6* | *AaHC7* | *AaHC8* | *AaFBC1* | *AaFBC2* | *AaFBC3* | *AaFBC4* | *AaFBC5* | *AaMuscle* |
|---|---|---|---|---|---|---|---|---|---|---|---|---|---|---|
| ***AgHC1*** | 77.49 | 99.51 | 6.92 | 1.27 | 2.08 | 2.44 | 0 | 0 | 3.73 | 1.99 | 7.20 | 0 | 0 | 6.52 |
| ***AgHC2*** | 10.21 | 0.49 | 50.38 | 17.20 | 25.00 | 13.01 | 2.67 | 3.49 | 16.00 | 9.93 | 34.09 | 0.72 | 9.68 | 43.48 |
| ***AgHC3*** | 6.54 | 0 | 19.25 | 53.82 | 19.79 | 66.67 | 4.00 | 1.16 | 0 | 2.32 | 3.79 | 0 | 0 | 2.17 |
| ***AgHC4*** | 0 | 0 | 4.81 | 0 | 0 | 2.44 | 86.67 | 0 | 0.27 | 0.33 | 0 | 0 | 0 | 0 |
| ***AgHC5*** | 0.79 | 0 | 3.61 | 3.82 | 19.79 | 0 | 0 | 1.16 | 0 | 0 | 0 | 0 | 0 | 0 |
| ***AgHC6*** | 1.57 | 0 | 7.97 | 6.69 | 8.33 | 6.50 | 4.00 | 94.19 | 1.07 | 0.33 | 0.38 | 0 | 0 | 0 |
| ***AgFBC2*** | 0 | 0 | 0.30 | 0 | 1.04 | 0 | 0 | 0 | 5.33 | 1.99 | 2.27 | 1.45 | 9.68 | 0 |
| ***AgFBC1*** | 3.40 | 0 | 5.71 | 15.61 | 23.96 | 8.13 | 1.33 | 0 | 73.33 | 82.45 | 51.14 | 97.83 | 80.65 | 15.22 |
| ***AgMuscle*** | 0 | 0 | 1.05 | 1.59 | 0 | 0.81 | 1.33 | 0 | 0.27 | 0.66 | 1.14 | 0 | 0 | 32.61 |

**Table S5**. Marker genes of *Aedes aegypti* cell clusters. P-value adj = P value adjusted for multiple testing. Avg logFC = average log fold change between cluster of interest and all other clusters. Pct.1 = percentage of cells in the cluster with detectable marker expression. Pct.2 = percentage of cells in all other clusters with detectable marker expression. Electronic annotation.

(see separate file)

**Table S6.** Granulocyte percentage of LL3-silenced mosquitoes injected with HDF. Data are representative of two independent experiments, with at least 8 individual mosquitoes each. Unpaired t-test was used for statistical analysis.

| | | Rep 1 | | Rep 2 | | Merge | |
|---|---|---|---|---|---|---|---|
| **Granulocytes** | | **% ± std** | **N** | **% ± std** | **N** | **% ± ste** | **N** |
| dsLacZ | -HDF | 1.58 ± 0.34 | 9 | 1.88 ± 0.44 | 13 | 1.76 ± 0.26 | 22 |
| | +HDF | 4.3 ± 1.29 | 8 | 10.74 ± 1.18 | 13 | 8.29 ± 1.09 | 21 |
| dsLL3 | -HDF | 2.6 ± 0.89 | 10 | 0.89 ± 0.16 | 12 | 1.69 ± 0.44 | 22 |
| | +HDF | 1.34 ± 0.37 | 10 | 1.38 ± 0.26 | 11 | 1.36 ± 0.22 | 21 |

**Table S7.** Oenocytoids percentage of LL3-silenced mosquitoes injected with HDF. Data are representative of two independent experiments, with at least 8 individual mosquitoes each. Unpaired t-test was used for statistical analysis.

| | | Rep 1 | | Rep 2 | | Merge | |
|---|---|---|---|---|---|---|---|
| **Oenocytoids** | | **% ± std** | **N** | **% ± std** | **N** | **% ± ste** | **N** |
| dsLacZ | -HDF | 19.7 ± 2.05 | 9 | 19.2 ± 2.15 | 13 | 19.4 ± 1.39 | 22 |
| | +HDF | 34.5 ± 2.21 | 8 | 43.5 ± 1.89 | 13 | 40.1 ± 1.69 | 21 |
| dsLL3 | -HDF | 19.3 ± 1.98 | 10 | 20.7 ± 2.33 | 12 | 20.1 ± 1.45 | 22 |
| | +HDF | 23.7 ± 1.99 | 10 | 22.3 ± 2.49 | 11 | 22.9 ± 1.61 | 21 |

**Table S8**. Prohemocytes percentage of LL3-silenced mosquitoes injected with HDF. Data are representative of two independent experiments, with at least 8 individual mosquitoes each. Unpaired t-test was used for statistical analysis.

| | | Rep 1 | | Rep 2 | | Merge | |
|---|---|---|---|---|---|---|---|
| **Prohemocytes** | | **% ± std** | **N** | **% ± std** | **N** | **% ± ste** | **N** |
| dsLacZ | -HDF | 78.7 ± 0.33 | 9 | 78.9 ± 2.10 | 13 | 78.8 ± 1.36 | 22 |
| | +HDF | 61.1 ± 3.16 | 8 | 45.7 ± 2.17 | 13 | 51.6 ± 2.43 | 21 |
| dsLL3 | -HDF | 78.0 ± 2.36 | 10 | 78.3 ± 2.36 | 12 | 78.2 ± 1.56 | 22 |
| | +HDF | 74.9 ± 2.03 | 10 | 76.3 ± 2.56 | 11 | 75.65 ± 1.65 | 21 |

**Table S9**. Summary of statistical analysis and P-values for extended tables S6, S7 and S8.

| Comparison | Granulocytes | Oenocytoids | Prohemocytes |
|---|---|---|---|
| dsLacZ -HDF Vs dsLacZ +HDF | ***, *P< 0.0001*, t= 5.897, df=41 | ***, *P< 0.0001*, t= 9.851, df=41 | ***, *P< 0.0001*, t= 9.458, df=41 |
| dsLL3 -HDF Vs dsLL3 +HDF | ns, P= 0.5146, t= 0.6575, df=41 | ns, P= 0.2679, t= 1.123, df=41 | ns, P= 0.1898, t= 1.333, df=41 |

**Table S10**. Percentage and total number of prohemocytes positive for either LRR8 or PPO4. Data are representative of 2 independent experiments.

| | Replicate 1 | | Replicate 2 | | Rep 1 + Rep 2 | |
|---|---|---|---|---|---|---|
| **Prohemocytes** | **%** | **# of cells** | **%** | **# of cells** | **%** | **# of cells** |
| $LRR8^+$ | 85.8 | 151 | 85.96 | 98 | 85.9 | 249 |
| $PPO4^+$ | 14.2 | 25 | 14.04 | 16 | 14.1 | 41 |

**Table S11**. Percentage and total number of each granulocyte population: granulocytes ($LRR8^H$ / $PPO4^L$), Megacytes ($LRR8^L$/ $PPO4^L$ neg/ $TM7318^+$) and Small granulocytes ($LRR8^L$/ $PPO4^L$/ $TM7318^-$). Data are representative of 2 independent experiments.

| | Replicate 1 | | Replicate 2 | | Rep 1 + Rep 2 | |
|---|---|---|---|---|---|---|
| **Granulocytes** | **%** | **# of cells** | **%** | **# of cells** | **%** | **# of cells** |
| $LRR8^H$ / $PPO4^L$ | 81.2 | 1017 | 85.2 | 1014 | 83.2 | 2031 |
| $LRR8^L$/ $PPO4^L$ neg/ $TM7318^+$ (megacytes) | 0.6 | 8 | 0.3 | 3 | 0.5 | 11 |
| $LRR8^L$/ $PPO4^L$/ $TM7318^-$ | 18.1 | 227 | 14.5 | 173 | 16.4 | 400 |

**Table S12**. Area-normalized number of granulocytes, oenocytoids, megacytes and AM granulocytes on the body of control and infected mosquitoes. Data are representative of 3 independent experiments. Welch T-Test was used for statistical analysis.

| | | Rep1 | | Rep2 | | Rep3 | | Merge | |
|---|---|---|---|---|---|---|---|---|---|
| | | **Total cells ± std** | **Welch T-Test** | **Total cells ± std** | **Welch T-Test** | **Total cells ± std** | **Welch T-Test** | **Total cells ± std** | **Welch T-Test** |
| **PPO$^{low}$/LLR8$^{high}$ granulocytes** | BF | 158.54 ± 50.04 | **, P = 0.0016 | 360.93 ± 72.10 | * , P = 0.0153 | 434.64 ± 110.86 | ** , P = 0.0056 | 285.68 ± 138.06 | **** , P < 0.0001 |
| | Pb | 50.13 ± 23.00 | | 150.22 ± 51.11 | | 122.99 ± 37.11 | | 89.56 ± 53.27 | |
| **PPO$^{high}$/LLR8$^{low}$ oenocytoids** | BF | 4.42 ± 4.75 | - , P = 0.0853 | 11.52 ± 6.70 | - , P = 0.9742 | 22.39 ± 20.94 | - , P = 0.2737 | 11.81 ± 14.73 | - , P = 0.4561 |
| | Pb | 0.42 ± 0.41 | | 11.26 ± 8.89 | | 8.75 ± 6.28 | | 5.91 ± 7.77 | |
| **TM7318+ megacytes** | BF | 1.93 ± 0.98 | - , P = 0.6284 | 2.57 ± 1.46 | - , P = 0.1747 | 2.80 ± 0.86 | - , P = 0.339 | 2.36 ± 1.16 | - , P = 0.7361 |
| | Pb | 1.49 ± 1.52 | | 1.08 ± 0.07 | | 3.80 ± 1.43 | | 2.16 ± 1.81 | |
| **TM7318- AM granulocytes** | BF | 0.46 ± 0.42 | - , P = 0.4785 | 0.51 ± 0.31 | - , P = 0.4112 | 5.96 ± 7.76 | - , P = 0.8251 | 2.19 ± 5.04 | - , P = 0.677 |
| | Pb | 0.69 ± 0.52 | | 1.16 ± 0.89 | | 6.97 ± 3.56 | | 2.90 ± 3.74 | |

**Table S13**. Surface area normalization of control and infected mosquito bodies. Data are representative of 3 independent experiments. Welch T-Test was used for statistical analysis. Area represented on supplementary table S12.

| | Rep1 | | Re2 | | Rep3 | | Merge | |
|---|---|---|---|---|---|---|---|---|
| | Area ± std (N) | Welch T-Test | Area ± std (N) | Welch T-Test | Area ± std (N) | Welch T-Test | Area ± std (N) | Welch T-Test |
| BF | 4.52 ± 0.71 (7) | - , P = 0.3234 | 4.78 ± 0.48 (4) | - , P = 0.6332 | 4.94 ± 0.45 (5) | - , P = 0.2695 | 4.72 ± 0.61 (16) | - , P = 0.6435 |
| Pb | 4.17 ± 0.33 (5) | | 4.96 ± 0.33 (3) | | 5.64 ± 0.87 (4) | | 4.86 ± 0.60 (12) | |

**Table S14**. Percentage and total number of granulocytes (LRR8+) expressing high and low levels of FBN29831 in control and infected mosquitoes. Data are representative of 2 independent experiments. Chi square test was used for statistical analysis.

| | | Rep 1 | | Rep 2 | | Merge | |
|---|---|---|---|---|---|---|---|
| Granulocytes LRR8H / PPO4L | | % | # of cells | % | # of cells | % | Total of cells |
| Control | FBN29831L | 99.0 | 98 | 65.1 | 456 | 82.0 | 800 |
| | FBN29831H | 1.0 | 1 | 34.9 | 245 | 18.0 | |
| Infected | FBN29831L | 6.5 | 10 | 40.9 | 610 | 23.7 | 1646 |
| | FBN29831H | 93.5 | 144 | 59.1 | 882 | 76.3 | |

**Table S15**. Chi square statistical analysis of granulocyte (LRR8+) proportions represented on supplementary table S14.

| | C Vs I - LRR FBN Adjusted Chi square |
|---|---|
| Chi-square, df | 747.7, 1 |
| z | 27.34 |
| P value | <0.0001 |
| P value summary | **** |

**Table S16**. Percentage and total number of oenocytoids ($PPO4^+$) expressing high and low levels of FBN29831 in control and infected mosquitoes. Data are representative of 2 independent experiments. Chi square test was used for statistical analysis.

| | | Rep 1 | | Rep 2 | | Merge | |
|---|---|---|---|---|---|---|---|
| **Oenocytoids $LRR8^L$ / $PPO4^H$** | | **%** | **# of cells** | **%** | **# of cells** | **%** | **Total of cells** |
| Control (C) | $FBN29831^L$ | 80 | 8 | 76.9 | 60 | 78.5 | 88 |
| | $FBN29831^H$ | 20 | 2 | 23.1 | 18 | 21.5 | |
| Infected (I) | $FBN29831^L$ | 9.4 | 3 | 58.1 | 90 | 34 | 186 |
| | $FBN29831^H$ | 90.6 | 29 | 41.3 | 64 | 66 | |

**Table S17**. Chi square statistical analysis of oenocyte ($PPO4^+$) proportions represented on supplementary table S16.

| | **C Vs I - PPO4 FBN Adjusted Chi square** |
|---|---|
| Chi-square, df | 47.46, 1 |
| z | 6.889 |
| P value | <0.0001 |
| P value summary | **** |

**Table S18**. FBN29831 gene expression in hemocytes from control and *P. berghei* infected mosquitoes 48 hours post feeding. Data are representative of two independent experiments; three biological replicates were used per condition for each experiment. Hemolymph of 20 mosquitoes was used for each biological replicate. Unpaired t-test was used for statistical analysis.

| Gene | Treatment | Rep 1 | Comparison | P value |
|---|---|---|---|---|
| FBN29831 | C | 1.047 ± 0.2224 | - | - |
| | I | 1.997 ± 0.04585 | C Vs I | p=0.0139 |

### Methods

#### *Anopheles gambiae* and *Aedes aegypti* rearing and *P. berghei* and *P. falciparum* infection

*Anopheles gambiae* (G3 NIH strain), *Anopheles gambiae* M-form (*A. coluzzii,* BEI resource number MRA-1279), and *Aedes aegypti* (Liverpool strain) were reared at 28°C, 80% humidity, 12 h light-dark cycle and maintained with 10% Karo syrup solution during adult stages (sugar fed mosquitoes). Infections with *Plasmodium berghei* were performed using transgenic GFP parasites (ANKA GFPcon 259cl2) kept by serial passages in 3 to 6-week-old female BALB/c mice (Charles River, Wilmington, MA, USA) from frozen stocks[1]. Parasitemia was assessed by light microscopy following methanol-fixed blood-smears stained with 10% Giemsa and air-dried. Four to five-day-old female mosquitoes were fed when mice reached 3-5% parasitemia. Age matched uninfected mice were used to feed blood-fed control groups. After feeding, control and infected mosquitoes were kept at 19°C, 80% humidity and 12h light-dark cycle until the day of hemocyte or tissue collection. To confirm infection intensity at least 10 mosquito midguts were dissected 5 days post blood-feeding and oocysts counted by fluorescence microscopy. NF54 (wild-type *P. falciparum*) was maintained in $O^+$ human erythrocytes with RPMI 1640 medium with 25 mM HEPES, 50 mg/l hypoxanthine, 25 mM $NaHCO_3$, and 10% (v/v) heat-inactivated type $O^+$ human serum supplementation (Interstate Blood Bank, Inc., Memphis, TN) at 37°C and with a gas mixture of 5% $O_2$, 5% $CO_2$, and balance $N_2$. *Plasmodium falciparum* infections were done by diluting to 0.1% gametocytemia mature NF54 gametocytes. Mosquitoes were then allowed to feed with an artificial membrane feeder. NF54 with human red blood cells to 45% haematocrit was placed in warmed to 37C water-jacketed glass membrane feeders and mosquitoes allowed to feed for 20 minutes. Fed mosquitoes were then incubated at 26°C and 80% humidity. Infection levels (oocyst numbers) were checked by first dissecting midguts in 1× PBS and then staining them in 0.1% mercurochrome ahead of compound microscope visualization.

#### Hemocyte collection

Hemocytes were collected by gradually injecting in the thorax of cold-anesthetized mosquitoes 10 μL of anti-coagulant media (2 μL at a time) composed of 60% Schneider's insect media, 30% citrate buffer, 10% heat-inactivated fetal bovine serum, final pH 7.0-7.4, sterilized through a 0.22 μm syringe filter. Firepolished and thin-wall single barrel TW150-6 borosilicate glass capillaries 152 mm long with 1.5 / 1.12 OD / ID in mm were prepared with a Narishige PC-10 needle puller (heater N.2 mode, heat level 24.8). The tip of the needle was carefully cut open with fine tweezers and a gentle incision made in the lower abdomen with sterile micro-forceps[2]. We built an oil-free anti-coagulant buffer injection system composed of a Tritech Research microINJECTOR system with a microinjector All-Digital Multi-pressure system (MINJ-D) controller, a precision N2 cylinder pressure regulator for gas pressure control (TREG-N2) fitted with BS341 cylinder fittings for use in the United Kingdom (TREG-BR580), and a brass straight-arm needle holder (MINJ-4). The regulator was set at 20 psi. Hemolymph was collected as above with a steady pressure of 1 psi during injection until 10 μL were collected per mosquito from the lower abdomen into non-stick Eppendorf tubes to prevent cell attachment. Cells were manually counted manually by placing 8-12 mosquito hemocytes in sterile single-use disposable hemocytometer slides (Neubauer Improved, iNCYTO C-Chip DHC-N01) under a light microscope with a 40X objective.

When preparing cells for scRNA-seq, hemocytes were treated with the biomolecule stabiliser and cell fixative *vivo*PHIX (RNAssist Ltd, Cambridge, UK). *vivo*PHIX protects RNA, DNA and proteins from degradation within fixed cell, was developed from a deep eutectic solvent, is non-cross-linking, dissolves fat droplets, and has very low volatility, so that fixed cells can be stored for weeks at room-temperature and months at 4°C prior to analysis by scRNA-seq. When fixing hemocytes with *vivo*PHIX cells were collected as above and then plunged into 500 μL of fixative at room temperature. After processing four mosquitoes the cell-fixative was mixed well by pipetting 5 times with a P1000. The procedure was repeated after adding four more samples, or reaching required amounts (8-12 mosquitoes per condition). Hemocytes were then fixed for 2 hours at room temperature, before being transferred to 4C storage. On the day of processing, fixed hemocytes were mixed with one volume of pure, molecular grade ethanol before centrifugation for 30 minutes at 3k RCF at room temperature. Supernatant was discarded and the pellet resuspended in pure molecular grade water before 10X Chromium scRNA-seq library processing.

***Anopheles gambiae* dsRNA micro-injections and LL3 knockdown**

Two to three-day old female *A. gambiae* G3 mosquitoes were cold anesthetized and injected with 69 nl of 3 µg/µl dsRNA solution specific for LacZ, a bacterial gene not found in the genome of mosquitoes. dsRNA of LacZ is used as control during dsRNA-injection gene knockdown. A 218-bp fragment was amplified from LacZ gene cloned into pCRII-TOPO vector using M13 primers to add a T7 tail[3]. For dsLL3 synthesis, a fragment was amplified with a T7 tail using the following primers: T7-LL3 F - TTAATACGACTCACTATAGGGAGAATGACTACCATCATAGTGACGAACCC and T7-LL3 R – TTAATACGACTCACTATAGGGAGATTACACCATTATTAAATAAATAACACAACTTGAG, as described before[4]. The PCR product, from LacZ and LL3, was used as a template for dsRNA synthesis with Megascript RNAi kit (ThermoFisher Scientific, Waltham, MA, USA) according to the manufacturer's instructions.

**Generation of Naïve (- HDF) and Challenged (+ HDF) and injection in LL3-knockdown mosquito recipients**

Mosquitoes were infected with *P. berghei* and following blood feeding, the naive group was placed at 28°C to prevent infection; while the challenged group was maintained at 21°C for 48h, for normal infection to proceed. Subsequently, the challenged group was transferred to 28°C to reduce parasite load. Hemolymph from naïve and challenged groups was collected at seven days post-infection and centrifuged at 4°C, 10,000 rpms for 10 min. The cell-free supernatant was transferred to a new microcentrifuge tube and stored at -80°C until its use. To evaluate the effects of LL3 depletion on the hemocyte's capacity to respond to HDF, 2-3-day old mosquitoes injected with dsRNA for LL3 or LacZ (control) were then injected with 138 nl of cell-free hemolymph from Naïve (- HDF) or Challenged (+ HDF) donors at 3 days post-silencing. Hemocyte differentiation was assessed in two independent experiments at four days post-hemolymph transfer.

**Hemocyte quantification**

In short, an anticoagulant solution (95% Schneider's Insect medium and 5% citrate buffer) was injected into the thorax of mosquitoes, and 10ul of the perfused retrieved by performing a fine incision between the last two abdominal segments. Hemocyte quantification was conducted by depositing the collected perfused in a sterile disposable hemocytometer slide (Neubauer Improved, iNCYTO C-Chip DHC-N01) and differentiating hemocyte populations under a light microscope (40X objective).

***Aedes aegypti* samples for scRNAseq**

Three different challenges were used to generate *A. aegypti* samples. Three to four days old females were either fed with serum, bacteria or injected with dsRNA for a bacterial gene (LacZ) and non-related to the mosquito genome. Mosquitoes were injected with dsLacZ two days prior to serum feeding. For the bacterial feeding we used a mixture of cultivable bacteria from our colony at NIH as described before[5]. Briefly, on the day of the experiment, a pre-inoculum was diluted in fresh LB and grown for 2 hours at 28°C. The culture was washed with PBS to remove toxins and the concentration was estimated by measuring the optical density (OD) at 600nm. We considered 1OD the equivalent of $10^9$ bacteria/ ml. Non-injected females were fed with either serum or $4x10^9$ bacteria mixed with serum. dsLacZ-injected females were fed only with serum. Four days post feeding, twenty-five mosquitoes were perfused per replicate and three replicates were done per experimental group (control serum fed, bacteria fed and dsRNA-injected serum fed). Hemolymph (10 µl/ mosquito) was placed directly in 0.5ml of *vivo*PHIX for stabilization. After 60 µm filtering three volumes of glacial acetic acid were added to one volume of fixed hemocyte and mixed well. After 10 minutes incubation samples were transferred to ice. Then, one volume of pure molecular grade ethanol was added to the mixture and mixed well before centrifugation for 20 minutes at 3k RCF at room temperature. Supernatant was discarded and pellet resuspended in pure molecular grade water with 0.1% BSA, freshly-prepared, before staining and sorting. Sony SH800 was used to sort *vivo*PHIX fixed hemocytes stained for 20 minutes with 1 drop per 500 µl of sample of NucBlue Live ReadyProbes Reagent (Hoechst 33342 formulation by ThermoFisher). The sorter was operated with 100 µm disposable chips. Cells were sorted on fluorescence intensity, with 405 nm laser excitation and Hoechst 33342 filter, gated to exclude negative events with non-stained control. Forward scatter (FSC) and side scatter (SSC) information was also used to exclude doublets and multiplets. Cells were sorted into chilled 1.5 mL Eppendorf tubes before scRNA-seq V2 3' Chromium 10X library preparation.

**Chromium 10X sc-RNAseq library preparation**

After having prepared an appropriate single cell suspension, 10X Genomics Chromium droplet single-cell RNAseq master mix was prepared and protocol followed per manufacturer's instructions for V2 3' Chromium 10X single cell kit (custom 14 PCR amplification cycles).

**Mosquito RNA extraction and bulk RNAseq library preparation**

For bulk RNAseq hemocytes were collected as described above from 8 mosquitoes, and transferred directly in 500 µL of TRIZOL reagent (Invitrogen). From the same mosquitoes, midguts and carcasses were transferred into separate 1.5 mL Eppendorf tubes containing 150 μL TRIZOL reagent. The samples were triturated with an electrical homogenizer and disposable pestles before adding 350 μL more TRIZOL reagent and mixing. Samples were lysed for 15-30 minutes at room temperature, then stored at 4°C overnight and then at -20 °C until RNA extraction. Non-hemocyte samples were spun for 12,000 RCF, 10 minutes at 4°C to remove all insoluble material. The supernatant, as well as the homogenate of hemocyte samples were transferred to Phase Lock GelHeavy (5PRIME) 2 mL tubes pre-spun at 1,500 RCF for 1 minute, and allowed to incubate for 5 minutes at room temperature. 100 μL of chloroform (200 μL per 1 mL TRIZOL) was added, the tubes capped, and then vigorously shaken for 15 seconds. Samples were centrifuged for 12,000 RCF, 10 minutes, 4°C. If the clear, aqueous phase was still mixed with TRIZOL matrix then 100 μL more of chloroform was added, and the samples again mixed vigorously and spun as before. The aqueous phase was transferred to a fresh 1.5 mL Eppendorf tube and the RNA precipitated by adding 0.25 mL of isopropyl alcohol (500 mL per 1 mL TRIZOL reagent used). For midguts and hemocyte samples 20 μL of glycogen (5 mg / mL) were also added to aid in precipitation and pelleting. Samples were mixed by repeated inversion 10 times, incubated at 10 min at room temperature, and spun at 12,000 RCF, 10 min, 4°C. The supernatant was removed, and the RNA pellets washed twice with 75% ethanol (minimum 1 mL of ethanol per 1 mL of TRIZOL used). Each time the samples were mixed by vortexing and centrifuged 7,500 RCF, 5 min, 4°C. Supernatant was removed and samples air-dried. RNA was resuspended with 20 μL of RNAse free water for hemocyte samples, 30 μL for midgut samples, and 70 μL for carcass samples, homogenized and then incubated at 55°C for 10 min. Samples were stored at -20°C until library preparation. Total RNA quantity was assessed on a Bioanalyser (300 ng to 39 μg). mRNA was isolated with the NEBNext Poly(A) mRNA magnetic isolation module (New England Biolabs) and RNA-seq libraries prepared using the NEBNext Ultra II Directional RNA Library Prep Kit for Illumina (New England Biolabs) as by manufacturer instructions, except that a proprietary Sanger UDI (Unique Dual Indexes) adapters / primer system was used and a Kapa Hifi polymerase rather than NEB Q5 was employed.

**Sequencing**

For bulk RNAseq samples HS4000, (using kit version 1) 75PE (RNA): libraries were run on the Illumina HiSeq 4000 instrument with standard protocols using a 150-cycle kit set to a 75bp paired-end configuration. Libraries supplied at 2.8 nM and loaded with a loading concentration of 280 pM. For scRNA-seq Chromium 10X V2 and V3 kits, HS4000 (using kit version 1) 10X V2 and V3 read lengths: libraries were run on the Illumina HiSeq 4000 instrument with standard protocols using a 150-cycle kit set. As recommended by 10x

Genomics an elongated reverse read was used during the sequencing run. For V2, the read lengths were as follows: Read 1: 26 bases, index 1: 8 bases, read 2: 98 bases. For V3, read lengths were as follows: Read 1: 28 bases, index 1: 8 bases, read 2: 91 bases. Libraries supplied at 2.8 nM and loaded at a concentration of 280 pM.

**Hemocyte RNA extraction, cDNA synthesis and qPCR analysis**

For qPCR, *A. gambiae* hemocytes were collected as described above 48 hours after control feeding and *P.berghei* infection. Hemolymph pools of 20 mosquitoes were placed directly into TRIzol LS reagent (ThermoFisher Scientific, Waltham, MA, USA). Two hundred microliters of chloroform were added (1/5 of TRIzol volume) to 1mL of TRIzol plus the hemolymph for the aqueous phase separation. Total RNA was precipitated with 2-propanol (1:1) added to the aqueous phase and 10 μg glycogen as a carrier. Total extracted RNA was resuspended in nuclease free water and one microgram was used for cDNA synthesis using the Quantitect reverse transcription kit (Qiagen, Germantown, MD, USA) following the company instructions. Quantitative PCR (qPCR) was used for measuring FBN29831(AGAP029831) gene expression in hemocytes cDNA. We used the DyNamo SYBR green qPCR kit (ThermoFisher Scientific, Waltham, MA, USA) with target specific primers and the assay ran on a CFX96 Real-Time PCR Detection System (Bio-Rad, Hercules, CA, USA). A 135-bp fragment was amplified for FBN29831 (F- ACTACCGAGACGAAAAGCCC and R- AACCACTAACCAACCTCCGC). Relative expression was normalized against *An. gambiae* ribosomal protein S7 (RpS7) as internal standard and analyzed using the ΔΔ Ct method[6,7]. RpS7 (AGAP010592) primers sequences were: F- AGAACCAGCAGACCACCATC and R – GCTGCAAACTTCGGCTATTC. Statistical analysis of the fold change was performed using Unpaired t-test (GraphPad, San Diego, CA, USA). Each independent experiment was performed with three biological replicates (three pools of 20 mosquitoes) for each condition.

**Hemocyte collection for morphology staining**

Hemocytes from *A. aegypti* (Liverpool) and *A. gambiae* (G3-CDC) were collected by perfusion, using anticoagulant buffer (60% Schneider medium, 30% citrate buffer (pH 4.5) and 10% FBS. PH was adjusted to 7 -7.2 after mixing all the components. After perfusion, hemocytes were placed in a μ-slide angiogenesis chamber (ibidi GmbH, Gräfelfing, Germany). Cells were fixed for an hour at room temperature by adding 16% paraformaldehyde (PFA) solution in anticoagulant buffer to a final concentration of 4%. After fixation cells were washed with PBS 0.1% Triton and incubated for 30 min at room temperature with 1U of phalloidin (Alexa Fluor 488 or 750, Molecular Probes, ThermoFisher Scientific, Waltham, MA, USA) and 20 µM Hoechst 33342 (405, Molecular Probes, ThermoFisher Scientific), both diluted in PBS. Cells were then placed in mounting media for storage by adding 2 drops of Prolong Gold Antifade Mountant (Molecular Probes, ThermoFisher Scientific).

**Morphology and *in situ* hybridization (ISH) analysis**

The ISH protocol includes permeabilization step with a protease and the morphology of the cell is completely lost after the treatment. In order to evaluate the morphology of hemocytes and RNA expression by FISH, we developed a two-step protocol to capture both events. Hemocytes collected by perfusion 48 h after control feeding or *P. berghei* infection were fixed and stained with Alexa 750 phalloidin (actin) as described above. Ten random fields of each well were imaged using a tile scan "mark and find" tool, where coordinates of the field are recorded and can be restored image the same cells later. Then, hemocytes were subjected to ISH using RNAscope multiplex fluorescent reagent kit v2 assay (cat# 323110, ACDBio, Abingdon, United Kingdom) according to manufacturer's instructions. TSA based fluorophores Opal 4- color automation IHC kit (cat # NEL801001KT, PerkinElmer, Waltham, MA, USA) was used for the development of fluorescence (Opal 520- C1, Opal 570 – C2; Opal 620 – C3 and Opal 690 – C4). Specific RNA probes were synthetized by ACDBio (Abingdon, United Kingdom) and are listed as follows: LRR8 (cat# 543211-C4; Aga-LRR-C4), PPO4 (cat# 543291-C2; Aga-PPO4-C2), TM7317 (cat# 543201-C3; Aga-Transmembrane-C3) and FBN29831 (cat# 543271-C1; Aga-Fibrinogen CT). At the end of the ISH protocol, hemocytes were placed in prolong gold and re-imaged using the "mark and find" tool to recall the positions of the morphology pictures. Images were merged using Imaris 9.3.1 (Bitplane, Concord, MA, USA).

**ISH quantitation analysis**

Images of 0.4 µm optical sections were acquired using a Leica TCS SP8 confocal microscope (Leica Microsystems, Wetzlar, Germany) with a 40x objective and tile scan mode. Images were processed and analyzed with Imaris 9.3.1 (Bitplane, Concord, MA, USA) using "spot function" tool for counting. First, we counted the number of cells in each field using Hoechst staining (nuclei), we applied a size threshold of 3 µm to avoid counting unspecific specks. To quantify FBN29831, TM7318, LRR8 and PPO4, we used a size threshold of 5 µm and applied a variable intensity sum filter depending on the channel. For FBN29831- lower threshold 2600 and higher threshold 1.46e4; for TM7318 – lower threshold 5077 and higher threshold 1.21e4; for LRR8 – lower threshold 1.04e4 and higher threshold 3.03e4; for PPO4 – lower threshold 2500 and higher threshold 3.02e4. Only cells that fell under those thresholds were counted and used for statistical analysis.

**Whole-mount FISH**

Mosquitoes were cold anesthetized, micro-injected with 69 nL of 16% fresh paraformaldehyde (PFA) and after 15 seconds dissected while bathing in freshly prepared 4% PFA. Carcasses and midguts were separated by adding carcasses directly into an Eppendorf containing 4% PFA on ice, while midguts were fixed for one minute in ice-cold fresh 4% PFA and then transferred to fresh 1X PBS where they were opened along their longitudinal axis with two small gauge needles under the dissecting microscope to release the blood meal. Using PBS surface tension all blood was released from the guts, which were then fixed in a 1.5 mL Eppendorf tube containing fresh 4% PFA. Samples were fixed overnight at 4°C on a rocker. Non-stick tubes and pipette tips were used to prevent sample adhesion. In all next steps care was shown in removing solutions, as guts especially can stick onto or be sucked into pipette tips, or remain stuck on tube walls. Solutions were always removed against a source of light to increase contrast and decrease likelihood to remove samples by error. Each wash was performed on a gentle rocker, as samples were fragile and could easily break apart.

The day after collection all PFA was removed and guts and carcasses washed twice with 1mL of PBST (0.1% v/v Tween 20 in 1x PBS). Hybridizations with RNAscope probes were carried out according to manufacturer's instructions, using RNAscope multiplex fluorescent reagent kit v2 assay (cat# 323110, ACDBio, Abingdon, United Kingdom) and nuclei stained with DAPI after the last wash. Following a final wash to remove excess DAPI, slides were mounted in a drop of Prolong Gold Antifade Mountant (Molecular Probes, ThermoFisher Scientific). The samples were flattened in the reagent under a dissecting microscope to prevent flaps and folding of the tissue. After adding coverslips, corners were fixed with transparent nail polish and the samples allowed to dry overnight at room temperature in the dark. The day after, nail polish was added all around the slide to seal the samples. These were then stored at 4°C in the dark until imaging.

**Mosquito sections FISH**

Mosquitoes were cold anesthetized, dipped in 100% ethanol to decrease surface tension, and then dipped and fixed in 10% formalin for 18-24 hours overnight at room temperature. The Sakura Tissue-Tek VIP Tissue processor on Rapid Biopsy programming was used (10 min VIP1 and 10 min VIP2 for each solution except: no VIP2 for 50% and 70% ethanol; first paraffin wax 20 min for both VIP1 and VIP2), with the following solutions in order: 50% ethanol, 70% ethanol, 90% ethanol, 3X 100% ethanol, 3X xylene, and 4x wax. For embedding, two orientations (longitudinal and transverse) were used for each condition (sugar-fed, blood-fed and *P. berghei* infection), before 5 μm sectioning. H&E sections were prepared for every other section, with the mirror section available for RNA-FISH (RNAscope, ACDBio) as of above.

**Imaging – slide scanning, confocal microscopy and tile scan imaging**

Mosquito sections and whole mounts were imaged with the 3DHISTECH MIDI II automatic digital slide scanner (3DHISTECH, Budapest, Hungary), with 20x and 40x objectives (numerical aperture 0.8 to 0.95), and a bespoke DAPI, Opal 520, 570, 620 and 690 filter sets and a 4.2MP 16-bit camera with wideband LED, or with a 20x bright-field camera for H&E mosquito sections and a 4.2MP 16-bit camera with RGB illumination. Sections and whole-mounts were imaged with extended focus, sequential acquisition, and variable z-steps, mosaic size and integration. For whole-mount and hemocytes samples images were captured at the NIH using a Leica TCS SP8 DMI8 confocal microscope (Leica Microsystems, Wetzlar, Germany) with a 20x, 40x and 63x oil immersion objective (using zoom factor of 2, 3 or 4; numerical aperture, 1.25 to 1.4) equipped with a photomultiplier tube/hybrid detector. Hemocytes were visualized with a white light laser, using 498-nm excitation for Alexa 488 (phalloidin) and Opal520; 550-nm excitation for Opal570; 588-nm excitation for Opal620; 670-nm excitation for Opal690 and a 405-nm diode laser for nuclei staining (Hoechst 33342). For Alexa 750nm, an Obis 690 laser was used for excitation and APD (avalanche photodiodes) detector was required for infrared detection. Images were taken using sequential mode and variable z-steps. For combined morphology and in RNA *in situ* hybridization, we used tile scan "mark and find" tool included in LASX software to capture the same areas of the slide before and after the hybridization. Image processing and merge was performed using Imaris 9.3.1 (Bitplane, Concord, MA, USA) and Adobe Photoshop CC (Adobe Systems, San Jose, CA, USA). At the Wellcome Sanger Institute images were captured using a Leica TCS SP8 DMI8 confocal microscope (Leica Microsystems) using a 40×, 63×, or 100× oil immersion objective (using zoom factor of 2, 3 or 4; numerical aperture, 1.25 to 1.4) and equipped with photomultiplier tube/hybrid detectors. Fluorochromes were excited using a 405 nm DMOD laser for DAPI, 488 nm CSU laser for Opal 520, a 552 nm CSU laser for Opal 570 and Opal 620, 638 nm CSU laser for Opal 690. Images were taken using sequential acquisition, and variable z-steps, mosaic size and integration. Image processing was performed using proprietary Leica LAS X and Imaris 9.2.1 (Bitplane, Concord, MA, USA). Whole-mount RNA-FISH positive cells were manually counted by an observer blinded to experimental conditions using the 3DHISTECH CaseViewer 2.3 software (3DHISTECH, Budapest, Hungary). Body wall and gut areas were measured with the analysis tools of the same software. For *P. falciparum* RNA-FISH experiments of hemocytes in circulation, positive cells were counted automatically using the segmentation and thresholding features of the Leica LAS X 3D visualization and analysis software (Leica UK, Milton Keys, UK). For *P. berghei* RNA-FISH experiments of hemocytes in circulation, image processing was performed using Imaris 9.3.1 (Bitplane, Concord, MA, USA).

**Transmission Electron Microscopy**

Two-days old female *A. gambiae* G3 mosquitoes kept in standard laboratory conditions were allowed to blood-fed once on an uninfected, anesthetized mouse. One day later they were anesthetized on ice, placed on a clean microscope slide on ice and decapitated with a sharp needle. Using fine forceps and iris scissors the dorsal body-wall was dissected free by means of longitudinal cuts along the left and right pleural regions, one transversal cut along A1 and another across A8. The tissue was flushed clean with cold Phosphate Buffered Saline (PBS) and immediately immersed in ice-cold fixative (see below) for 5h. The sample contained the dorsal cuticle with attached muscles, heart, abdominal fat body, tracheae and hemocytes. The fixative was a freshly prepared solution containing 2.5% glutaraldehyde, 4% paraformaldehyde and 2% tannic acid in 0.1M PBS pH 7.3. Fixation was stopped with 3x 10min washes in PBS. Thereafter the tissues were post-fixed for 1h in the darkness in an aqueous 1% solution of osmium tetroxide, washed a few times with H2O and dehydrated in an ascending series of ethanol (10-15 min each step). After dehydration the tissues were incubated 2x 15 min in propylene oxide (PO) and infiltrated gradually in epoxy resin (EPON 812 embedding kit 3132, Tousimis). Thereafter the samples were placed in new resin in a silicone rubber mould, oriented to obtain transverse sections and left 24h in an oven for resin polymerization at 60°C. Semithin serial sections (1.5 µm) were cut with a glass knife across abdominal segment A3 or A4, stained with boracic toluidine blue and analyzed by light microscopy to localize appropriate places for ultrastructural analysis. Three blocks (each containing one specimen) were retrimmed, the orientation was re-adjusted and ultrathin sections (50-60 nm) were obtained with a diamond knife. From each sample 20-30 sections were collected on clean, non-coated 300-mesh copper grids, contrasted with lead citrate and uranyl acetate and examined with a JEOL 100 CX electron microscope operated at 70 kV.

**Bulk RNA-seq analysis**

Sequencing reads in CRAM format were fed into a bespoke BASH pipeline to first automatically convert cram files to fastq using biobam's bamtofastq program (Version 0.0.191). Then, forward and reverse fastq reads in paired mode were aligned to the *Anopheles gambiae* AgamP4.9 reference genome using hisat2 (Version 2.0.4) and featureCounts (Version 1.5.1) with recommended settings. Combined counts matrix was then produced by a python script before downstream data processing and analysis within R version 3.5.3 (RStudio version 1.0.153). Downstream normalization, differential expression analysis and visualization were done with DESeq2 R package (Version 1.18.1)[8]. Base factor was defined as the sugar condition, and time 0 (non-infected). One outlier was removed (blood fed hemocyte sample at 48 hours) after plotting residuals of internal batch correction and visually inspecting a PCA plot. Data was normalized by making a scaling factor for each sample. First the log(e) of all the expression values were taken, then all rows (genes) were averaged (geometric average). Genes with zero counts in one or more samples were filtered out and the average log value from log(counts) for all genes was subtracted. Finally, the median of the ratios calculated as above for each sample was computed and raised to the e to make the scaling factor. Original read counts were divided by the scaling factor for each sample to get normalized counts. Then, the dispersion for each gene was estimated, and a negative binomial generalized linear model fitted. P-values for the differential expression analysis were adjusted for multiple testing using the Bonferroni correction. Genes were considered as differentially expressed if they had an adjusted P value $< 0.001$ (Wald T-test) and a $\log_2$ fold change $> 2$. All body parts, conditions and timepoints were considered together while running the following model for differential expression analysis focused on body part, with experimental repeats, time, and effects of treatment (*P. berghei*, blood feeding and sugar feeding) as covariates:

```
ddsMat <- DESeqDataSetFromMatrix(countData = countdata, colData = coldata,
            design = ~ 0 + experiment + time + treatment + part)
```

**scRNA-seq analysis**

Droplet-based sequencing data were aligned and quantified using the Cell Ranger Single-Cell Software Suite[9] (version 2.0, 10x Genomics) against the *Anopheles gambiae* PEST, AgamP4.9 reference genome provided by Vectorbase[10]. Cells with fewer than 100 and more than 2500 genes and for which total mitochondrial gene expression exceeded 20% (or 50%) were removed. Genes that were expressed in fewer than three cells were also removed. Downstream analyses—such as normalization, shared nearest neighbor graph-based clustering, differential expression analysis and visualization—were performed using the R package Seurat (3.0.2)[11-13]. The two experimental batches were integrated with a hybrid CCA / MNN strategy identifying 'anchors' of similar cells between conditions and CCs. Cells for which the expression profile could not be explained by low-dimensional canonical correlation analysis compared to low-dimensional principal component analysis were discarded. Clusters were identified using the community identification algorithm as implemented in the Seurat 'FindClusters' function. The resolution parameter to obtain the resulting number of clusters was fine-tuned so that it produced a number of clusters large enough to capture most of the biological variability. UMAP analysis was performed using the RunUMAP function with default parameters. Differential expression analysis was performed based on the Wilcoxon rank-sum test. The P values were adjusted for multiple testing using the Bonferroni correction. Clusters were annotated using canonical cell-type markers. We removed a blood-fed 24 hours post-feeding sample because it formed a technical outlier in the initial PCA-driven quality control. Hemocyte clusters were further analyzed by partitioning the clusters separately and performing the analysis anew, with the same alignment and clustering procedure.

Diffusion pseudotime[14] implemented in the SCANPY package[15] was applied to find the major non-linear components of variation across cells, using the most highly variable genes. Genes which changed along the identified trajectories (diffusion components) were identified by performing a likelihood ratio test using the function differentialGeneTest in the monocle 2 package[16].

Lineage tree reconstruction was performed with partition-based graph abstraction (PAGA) as implemented in SCANPY package[17]. The graph abstraction algorithm combines clustering and trajectory inference to elucidate the variability of scRNA-seq through discrete and continuous variables. PAGA takes into consideration a partitioned graph of neighborhood relations. It quantifies distances between nodes with a random-walk based measure and then it quantifies what connectivity partitions there is. The abstracted graph is anchored on nodes which are the clusters first identified with Seurat. The differentiation tree is a tree-like subgraph which best explains topology. Slingshot was another highly rated lineage tree reconstruction software that we used to validate PAGA results[18]. With a matrix input representing cells in a reduced-dimensional space (UMAP) and a vector of cluster labels, the Slingshot algorithms then built a minimum spanning tree (MST) of the clusters to infer the lineage structure. Finally, smooth lineage curves were built and pseudotime inferred for all lineages.

To compare the *A. gambiae* with the *Aedes* cell types, a logistic regression with L2-norm regularization and a multinomial learning approach (implemented by the scikit-learn function LogisticRegression) was trained on the anopheles gambiae clusters. The log-transformed normalized data was used. The model was used to predict the probabilities of each *Aedes* cell belonging to each one of the anopheles gambiae clusters (implemented by the predict_log_proba function).

**Additional references**


1 Franke-Fayard, B. *et al.* A Plasmodium berghei reference line that constitutively expresses GFP at a high level throughout the complete life cycle. *Mol Biochem Parasitol* **137**, 23-33, doi:10.1016/j.molbiopara.2004.04.007 (2004).

2 Rodrigues, J., Brayner, F. A., Alves, L. C., Dixit, R. & Barillas-Mury, C. Hemocyte differentiation mediates innate immune memory in Anopheles gambiae mosquitoes. *Science* **329**, 1353-1355, doi:10.1126/science.1190689 (2010).

3 Molina-Cruz, A. *et al.* Some strains of Plasmodium falciparum, a human malaria parasite, evade the complement-like system of Anopheles gambiae mosquitoes. *Proc Natl Acad Sci U S A* **109**, E1957-1962, doi:10.1073/pnas.1121183109 (2012).

4 Smith, R. C., Barillas-Mury, C. & Jacobs-Lorena, M. Hemocyte differentiation mediates the mosquito late-phase immune response against Plasmodium in Anopheles gambiae. *Proc Natl Acad Sci U S A* **112**, E3412-3420, doi:10.1073/pnas.1420078112 (2015).

5 Barletta, A. B. F., Trisnadi, N., Ramirez, J. L. & Barillas-Mury, C. Mosquito Midgut Prostaglandin Release Establishes Systemic Immune Priming. *iScience* **19**, 54-62, doi:10.1016/j.isci.2019.07.012 (2019).

6 Livak, K. J. & Schmittgen, T. D. Analysis of relative gene expression data using real-time quantitative PCR and the 2(-Delta Delta C(T)) Method. *Methods* **25**, 402-408, doi:10.1006/meth.2001.1262 (2001).

7 Pfaffl, M. W. A new mathematical model for relative quantification in real-time RT-PCR. *Nucleic Acids Res* **29**, e45, doi:10.1093/nar/29.9.e45 (2001).

8 Love, M. I., Huber, W. & Anders, S. Moderated estimation of fold change and dispersion for RNA-seq data with DESeq2. *Genome Biol* **15**, 550, doi:10.1186/s13059-014-0550-8 (2014).

9 Zheng, G. X. *et al.* Massively parallel digital transcriptional profiling of single cells. *Nat Commun* **8**, 14049, doi:10.1038/ncomms14049 (2017).

10 Giraldo-Calderon, G. I. *et al.* VectorBase: an updated bioinformatics resource for invertebrate vectors and other organisms related with human diseases. *Nucleic Acids Res* **43**, D707-713, doi:10.1093/nar/gku1117 (2015).

11 Satija, R., Farrell, J. A., Gennert, D., Schier, A. F. & Regev, A. Spatial reconstruction of single-cell gene expression data. *Nat Biotechnol* **33**, 495-502, doi:10.1038/nbt.3192 (2015).

12 Stuart, T. *et al.* Comprehensive Integration of Single-Cell Data. *Cell* **177**, 1888-1902 e1821, doi:10.1016/j.cell.2019.05.031 (2019).

13 Butler, A., Hoffman, P., Smibert, P., Papalexi, E. & Satija, R. Integrating single-cell transcriptomic data across different conditions, technologies, and species. *Nat Biotechnol* **36**, 411-420, doi:10.1038/nbt.4096 (2018).

14 Haghverdi, L., Buttner, M., Wolf, F. A., Buettner, F. & Theis, F. J. Diffusion pseudotime robustly reconstructs lineage branching. *Nat Methods* **13**, 845-848, doi:10.1038/nmeth.3971 (2016).

15 Wolf, F. A., Angerer, P. & Theis, F. J. SCANPY: large-scale single-cell gene expression data analysis. *Genome Biol* **19**, 15, doi:10.1186/s13059-017-1382-0 (2018).

16 Qiu, X. *et al.* Reversed graph embedding resolves complex single-cell trajectories. *Nat Methods* **14**, 979-982, doi:10.1038/nmeth.4402 (2017).

17 Wolf, F. A. *et al.* PAGA: graph abstraction reconciles clustering with trajectory inference through a topology preserving map of single cells. *Genome Biol* **20**, 59, doi:10.1186/s13059-019-1663-x (2019).

18 Street, K. *et al.* Slingshot: cell lineage and pseudotime inference for single-cell transcriptomics. *BMC Genomics* **19**, 477, doi:10.1186/s12864-018-4772-0 (2018).
